# Pax9 governs anterior identity and deployment of sclerotome to the median fins

**DOI:** 10.64898/2026.06.25.733239

**Authors:** Sarah McLeod, Margaret Keating, Ziyu Dong, Raisa Bailon-Zambrano, Katrinka M. Kocha, Sandhya Paudel, Abigail Mumme-Monheit, Melissa Scott-Preusse, Colette A. Hopkins, Raelyn Begay, Peng Huang, GuangJun Zhang, James T. Nichols, Lindsey Barske

## Abstract

The caudal fin is an anomaly among vertebrate locomotory appendages: its internal skeleton is as asymmetric as the human hand, but it lacks the Shh-secreting Zone of Polarizing Activity (ZPA) and the Gli3/HoxD/Hand2 programs that govern anterior-posterior patterning in other appendages. As the caudal fin was also the first appendage to evolve, deciphering its alternative patterning program may provide clues to the ancestral state. Pax9 is one of few conserved appendage patterning factors reported to also be active in the caudal fin, specifically in the anterior domain. We report that loss of *pax9* function in zebrafish not only disrupts anterior-specific caudal fin anatomy, but also results in a spectacular fusion of the caudal and anal fins along the ventral midline. The dorsal fin is also expanded to a lesser degree, the paired fins not at all. The mutant caudal fin initially forms as an irregularly patterned structure lacking anterior molecular identity, with supernumerary elements spilling out beyond the normal anterior boundary. Unexpectedly, this phenotype is subsequently compounded by neighboring trunk somites erroneously deploying skeletal mesenchyme in the normally finless caudal peduncle region, completing the ectopic skeleton. scRNAseq analysis at caudal fin bud stage indicates that *pax9* mutants gain skeletal mesenchyme at the expense of a specialized type of fibroblast involved in the formation of fin fold actinotrichia. Median fin skeletal mesenchyme and fin fold fibroblasts both arise from the sclerotome, a somite compartment that also robustly expresses *pax9*. We propose that, within the sclerotome, Pax9 pushes progenitors towards fin fold fibroblast fate, limiting how many cells will later be available to make median fin skeleton. Within the fin bud, it drives anterior identity, with the strongest impact on the ZPA-free caudal fin bud. These dual sites of action make Pax9 a uniquely powerful governor of median fin development.

## INTRODUCTION

The caudal fin serves the primitive, quintessential function of propelling a fish forward through the water. Its internal support skeleton directly attaches to the most posterior ural and preural vertebrae (Bird & Mabee, 2003), efficiently conveying the thrust forces generated by lateral undulation of the body through the tip of the tail (Webb, 1984). The other median fins (dorsal and anal) are not directly connected to the vertebral column and instead mainly serve to stabilize body posture against shifting water currents and forces produced by the other fins (Bird & Mabee, 2003). The paired pectoral and pelvic fins facilitate more complex maneuvers (Webb, 1984) and are the evolutionary predecessors of tetrapod fore- and hindlimbs (Paco & Freitas, 2018). Consistent with its cardinal function, the caudal fin emerges first in the fossil record, appearing in agnathans more than 430 million years ago (Soehn & Wilson, 1990; Larouche et al., 2019), well before the other median and paired appendages (Coates, 1994; Shu et al., 1999; Zhang & Hou, 2004). The prevailing hypothesis in the field is that developmental genetic programs for making appendages first evolved in the midline and were later co-opted to build the pectoral and pelvic appendages (Mabee et al., 2002; Freitas et al., 2006). This would have involved a shift from paraxial mesoderm – the source of the median fins (Freitas et al., 2006; Lee et al., 2013b; Shimada et al., 2013) – to lateral plate mesoderm, the source of the paired fins (Freitas et al., 2006; Prummel et al., 2020). The caudal fin, as the most ancient, paraxial mesoderm-derived median appendage, might thus best reflect the primitive state.

Though the adult caudal fin has served as an informative regeneration model for the past 30 years (Sehring & Weidinger, 2022), the programs controlling its development and morphology are not well understood in any species. Caudal fins that appear externally asymmetric (heterocercal) or symmetric (homocercal) both have an asymmetric endoskeleton. In the homocercal forked fin of the zebrafish, distinctly shaped endochondral support elements protrude from the ventral edge of the notochord (Bird & Mabee, 2003): in order from posterior to anterior, they are hypural elements 5, 4, 3, 2, 1, followed by the parhypural and two modified hemal spines of the preural vertebrae (Fig. 1A). The central diastema is located between hypurals 2 and 3. Exoskeletal fin rays, or lepidotrichia, are attached to the distal ends of each endochondral element and form the bulk of the external fin. As these elements are differentiating (<14 days post-fertilization (dpf) or 4-5 mm standard length (SL)), the tip of the notochord flexes dorsally, carrying the posterior elements upwards and aligning the diastema with the primary body axis to establish dorsoventral symmetry (Bird & Mabee, 2003; Cumplido et al., 2024). Consistent with this shape, the later stages of development and regeneration of the fin rays point to the existence of a central molecular organizer at the diastema, flanked by peripheral organizers at the anterior (later ventral) and posterior (later dorsal) boundaries (Desvignes et al., 2022). However, how the underlying asymmetric endoskeleton is patterned is unknown.

**Fig. 1.**
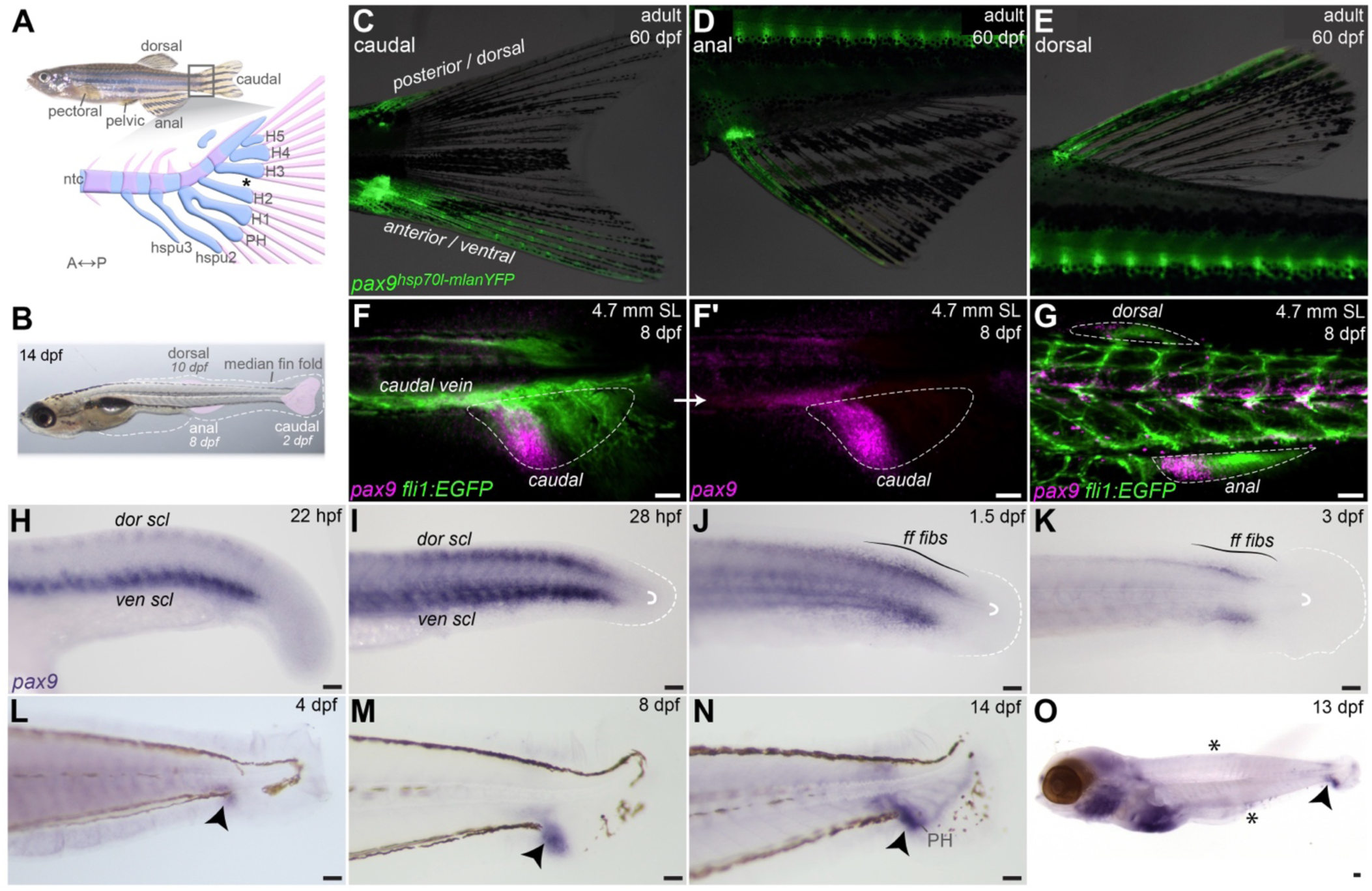
*pax9* is enriched at the anterior edges of all median fin fields. **A**, Adult zebrafish with fins marked and schematic marking key elements of caudal fin endoskeleton. The asterisk marks the diastema. H1-5, hypurals 1-5; PH, parhypural; hspu2-3, hemal spines of preural vertebrae; ntc, notochord. Blue elements are cartilaginous or notochord; pink elements are ossified bone. **B**, Location and approximate timing of appearance of the three median fin buds (pink) superimposed on a juvenile zebrafish. The caudal fin is already well developed at this stage. Dashed line outlines the median fin fold. **C-E**, A new *pax9* knockin reporter (*pax9^hsp70l:mlanYFP^*) strongly marks the anterior domains of each median fin (caudal (B), anal (C), dorsal (D)). A weaker domain is apparent along the posterior/dorsal edge of the caudal fin. Also see Fig. S1. **F-G**, *pax9* HCR (magenta) reveals anterior bud-restricted transcription in 8 dpf *fli1:*EGFP+ fish. Median fin buds are outlined. F’ shows the HCR channel alone. Images are maximum intensity projections. **H-O**, Colorimetric in situs for *pax9*. Earlier stages (H-J) reveal *pax9* expression in the dorsal and ventral sclerotome compartments (dor scl, ven scl) and fin fold fibroblasts (ff fibs). Expression is highly enriched in the nascent caudal bud (arrowheads) from 4 dpf on (L-N). Note enriched expression anterior to the parhypural at 14 dpf (N) and higher expression in the caudal relative to dorsal and anal fin buds at 13 dpf (O; asterisks, undetectable with this method). White semicircles and dashed lines in I-K outline the caudal termini of the notochord and fin fold, respectively. Scale bars F-O = 50 μm.

Like the paired appendages, the caudal and other median fins form as aggregations of mesenchymal cells at specific axial positions, with the zebrafish caudal bud appearing first at 2 dpf and the anal and dorsal buds at ∼7-8 and 10 dpf, respectively (Fig. 1B). These midline buds do not protrude externally from the body but instead develop inside the median fin fold, a transient structure comprised of epidermis, periderm, and fin fold fibroblasts that forms at 18 hpf then diminishes at juvenile stages when the median fin skeleton has differentiated (Abe et al., 2007; Miyamoto et al., 2022). Unlike all other fin and limb buds, the caudal fin bud lacks a Sonic Hedgehog (Shh)-secreting Zone of Polarizing Activity (ZPA), the organizer that imposes anterior-posterior polarity in other appendages (Hadzhiev et al., 2007). Whether this is an ancestral or derived condition in cyprinids is uncertain due to incomplete reporting in other taxa. In a bud with a ZPA, activation of Shh signaling is high in the posterior and low in the anterior, with the Gli3 effector accordingly stabilized in its full-length form in the posterior but processed into its truncated repressor form (Gli3R) in the anterior (Wang et al., 2000; Litingtung et al., 2002; te Welscher et al., 2002). Activation of Shh in posterior bud mesenchyme is induced in part by the Hand2 transcription factor (Charite et al., 2000; Galli et al., 2010). However, the zebrafish caudal fin bud does not express Hand2, Shh, or any other Hh ligand (Thisse & Thisse, 2005; Hadzhiev et al., 2007; Nachtrab et al., 2013; Tanaka et al., 2024). Mutants for *shha* and *smo* (Smoothened) present early axial defects that preclude bud initiation entirely (Hadzhiev et al., 2007), but the pathway is thereafter not active nor required for anterior-posterior caudal fin patterning (Braunstein et al., 2021). Curiously, other Hh pathway components (e.g. *gli3*, *gli2a*, *smo*) are nevertheless enriched in early caudal fin bud mesenchyme (Hadzhiev et al., 2007). Accordingly, the bud is sensitive to repatterning by ectopic Shh misexpression at 2 or 3 dpf, giving rise to a truncated rather than forked tail fin (Surette et al., 2025). There is no evidence of a requirement for *gli3* either: in medaka, *gli3* crispants developed supernumerary elements in the pectoral, pelvic, and dorsal fins (similar to mice and humans (Vortkamp et al., 1991; Hui & Joyner, 1993) but had no reported caudal (or anal) fin anomalies (Letelier et al., 2021). Zebrafish *gli3* mutants had no fin phenotypes at all (Tanaka et al., 2024).

Beyond lacking a ZPA, the caudal fin also lacks nested *Hoxd9-13* expression (Freitas et al., 2006), a complementary mechanism to the ZPA for determining anterior-posterior polarity in paired appendages (Zakany et al., 2004; Tarchini et al., 2006). Hox13 paralogs do pattern the caudal region of the fish: loss of *hoxc13a* or *hoxb13a* alone causes subtle abnormalities in ural/preural structures and the caudal fin endoskeleton, while their combined knockout results in loss of the caudal fin altogether due to homeotic transformation of segmental identity (Cumplido et al., 2024). Other combinations, e.g. *hoxa13a; a13b;d13a*, only reduce caudal fin length (Corcoran et al., 2026).

In contrast to the missing posteriorizing signals, several conserved anterior-restricted transcription factors (*pax9*, *alx4a*, *tbx3a*) are reportedly active in the anterior caudal fins of different fishes (Nachtrab et al., 2013; Schartl et al., 2021; Desvignes et al., 2022). Limited information is available on their functional requirement in this appendage, other than mild hypural abnormalities in medaka with transient morpholino knockdown of *pax9* (Mise et al., 2008). Pax9 is of special interest because, in addition to appendages, it is also expressed in the sclerotomal compartment of the paraxial mesoderm (Mise et al., 2008; Ma et al., 2018), from which the median fin skeleton originates (Freitas et al., 2006; Bailon-Zambrano et al., 2024). Homozygous loss of *Pax9* in mice results in preaxial polydactyly (an extra thumb), associated with expanded mesenchyme (Peters et al., 1998). Combined loss of both *Pax9* and its paralog *Pax1* causes major vertebral defects without worsening limb phenotypes (Peters et al., 1999), indicating that, at least in mice, *Pax9* is active in the sclerotome and in lateral plate mesoderm-derived structures, and redundant with *Pax1* only in the former.

In this work, we investigated the functional role for Pax9 in zebrafish caudal fin patterning and development. We uncover a unique dual requirement: not only does Pax9 establish and enforce the anterior boundary of the zebrafish caudal fin bud, but it also acts earlier within the sclerotome to influence how many cells contribute to the median fin skeleton *in toto*. Our results inspire a new model for how the caudal fin bud acquires anterior-posterior pattern in the absence of a ZPA or nested HoxD expression, a mechanism that might reflect the primitive appendage state.

## RESULTS

### pax9 robustly marks the anterior edge of all fin fields

To expand on the limited qRT-PCR and in situ data currently placing *pax9* in the caudal fin of adult medaka and swordtail fishes (Mise et al., 2008; Schartl et al., 2021), we employed a new zebrafish *pax9* YFP knock-in allele (*pax9^hsp70l-mlanYFP^ ^pu115^*, Fig. S1A). Clear YFP signal was evident along the anterior edge of all median fins in juvenile and adult zebrafish (Fig. 1C-E, S1B-G). As previously observed for another anterior factor, *alx4a* (Desvignes et al., 2022), a second, weaker domain also appears along the dorsal edge of the caudal fin after notochord dorsiflexion has instituted dorsoventral symmetry. We then evaluated endogenous *pax9* transcription at juvenile and larval stages using fluorescent hybridization chain reaction (HCR) assays and colorimetric in situs (Fig. 1F-O). HCRs were performed on fish carrying the *fli1:EGFP* transgene, which marks all fin bud mesenchyme as well as endothelial cells (Lawson & Weinstein, 2002; Bailon-Zambrano et al., 2024). By HCR, *pax9* was detected in the anterior compartment of all median fin buds (Fig. 1F-G) though more robustly in the caudal fin than the dorsal and anal fins. The less sensitive colorimetric alkaline phosphatase method detected only the caudal fin domain (Fig. 1O). At 14 dpf (∼5 mm SL), when endochondral elements have differentiated, the most intense staining was located anterior to the parhypural (Fig. 1N). At 4-8 dpf, prior to skeletal differentiation, *pax9* and *pax9^hsp70l-mlanYFP^* expression was enriched in the anterior part of the caudal bud, extending into the ventral part of the two somites directly dorsal to it (Fig. 1L-M, S1B).

Of interest, examination of earlier stages indicated that the latter caudal somitic domain may be a holdover from earlier *pax9* expression in the sclerotome. Zebrafish have a bipartite sclerotome, with distinct dorsal and ventral domains (Ma et al., 2018). As reported (Mise et al., 2008; Ma et al., 2018), *pax9* is expressed throughout both subcompartments beginning around 18 hpf (22 hpf shown in Fig. 1H). By 28 hpf, expression has declined in anterior somites relative to more recently formed posterior somites (Fig. 1I), remaining strong in vertebral precursors (Fig. 1G) and the two most caudal well-formed somites (Fig. 1J-K). The most posterior, unsegmented paraxial mesoderm located above the posterior part of the caudal bud and surrounding the notochord terminus does not express *pax9* (Fig. 1H-M).

At 1.5 and 3 dpf, *pax9* transcripts were also detected in mesenchymal cells out in the median fin fold (Fig. 1J-K). These appeared to be the sclerotome-derived fin fold fibroblasts that exit the somites early in development (26-30 hpf) and may help make the actinotrichia structures that support the fin fold (Zhang et al., 2010; Lee et al., 2013a). Reanalysis of the *pax9* knock-in line and published scRNAseq data confirm that this fibroblast cell population expresses *pax9* (Fig. S1B) (Sur et al., 2023).

### pax9 is an essential regulator of median fin development

To determine the requirement for *pax9* in caudal fin development, we used our *pax9^el622^* allele, which is a 10-bp deletion in exon 2 that leads to a frameshift at amino acid 34 of 259 and premature termination after 16 erroneous residues (Paudel et al., 2025). Unexpectedly, in adult homozygous mutants, the caudal and anal fins are expanded and fused along the ventral midline (Fig. 2A’, S2E, Movie S1). The unaffected dorsal (initially posterior) lobe of the caudal fin is longer than the ventral (initially anterior) lobe below the diastema (Fig. 2A’, S2E), resulting in heterocercal-like asymmetry. Fin pigmentation is irregular and variable between individuals (Fig. 2A’, S2C-E), with normal patterns observed only in the dorsal lobe of the caudal fin. Approximately 30% of mutants show weaker expressivity, with a small discontinuity interrupting the ectopic fin (Fig. S2C-D,O). However, in these cases, the caudal fin remains asymmetric, and the anal fin is still posteriorly elongated along the body axis. Conversely, the dorsal, pectoral, and pelvic fins were not overtly altered. Identical phenotypes were observed in homozygotes carrying a second, independently generated deletion allele also made on the Tübingen genetic background, *ci3038* (Fig. S2F-G) (Paudel et al., 2025). Three additional alleles, *pu116*, *ca122* and *ca123*, generated on a mixed TAB (AB/Tübingen) (*pu116*) or TL (Tüpfel longfin; *ca122* and *ca123*) background, rarely produced full fusion but consistently had asymmetric caudal fins and expanded anal fins (Fig. S2H-L) (mutation details for all alleles shown in Fig. S2A). The variable expressivity might be attributable to genetic background (Keating et al., 2026). Homozygotes for all five alleles are viable and fertile, and swim without apparent defects (Movie S1). *el622* was used for subsequent experiments.

**Fig. 2.**
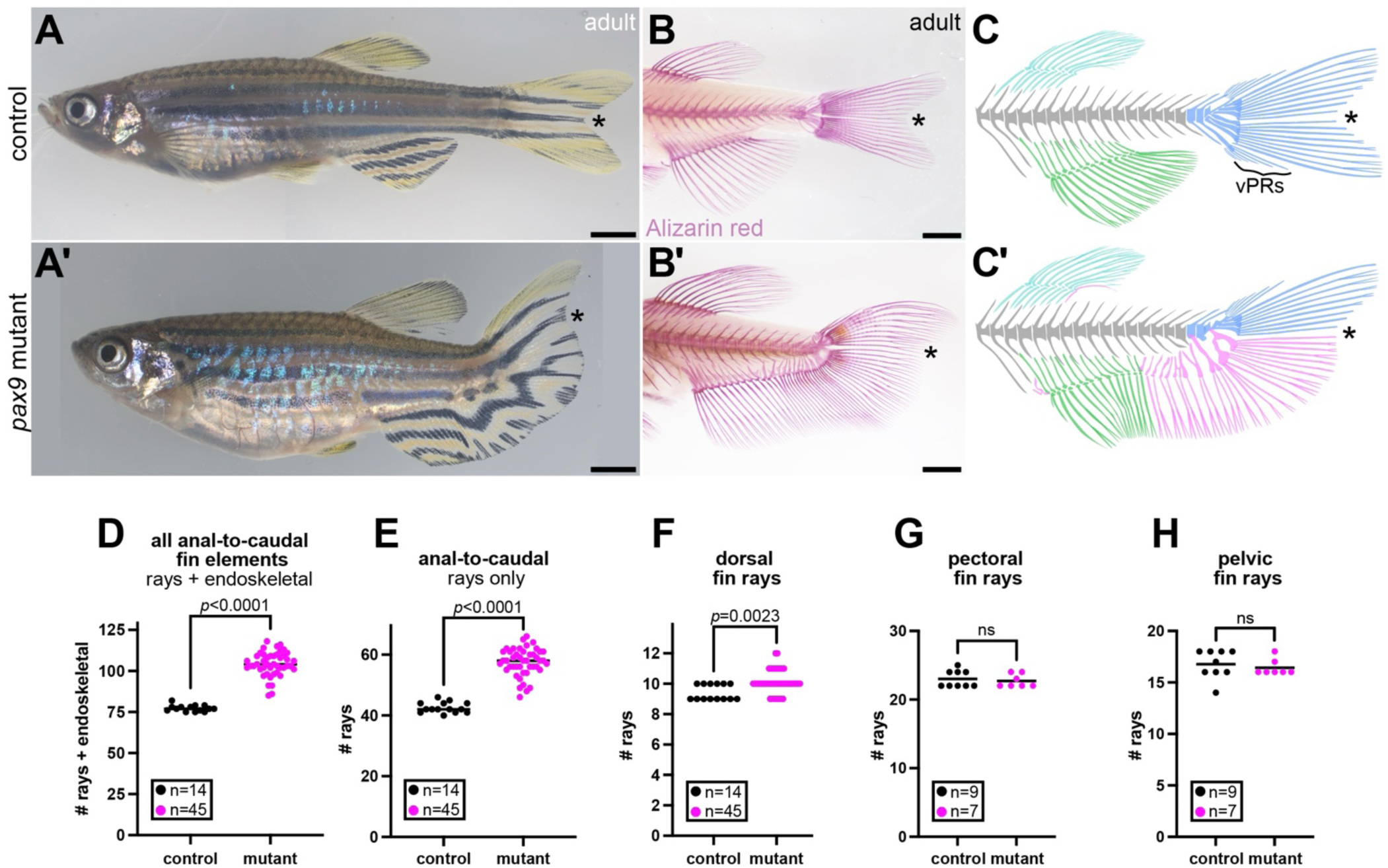
Pax9 is an essential regulator of median fin development. **A**, Brightfield images of adult control (A) and *pax9^el622^* mutant (A’) showing the continuous anal-to-caudal ventral fin. Note the merged pigmentation pattern. **B**, The mutant ectopic fin is fully supported by Alizarin red-stained mineralized bone. **C**, Schematics of median fin skeleton in the control and mutant condition, with deformed and ectopic elements highlighted in pink. vPRs, ventral procurrent rays. Scale bars in A-B = 2.5 mm. Asterisks mark the position of the diastema in A-C. **D-H**, Quantification of fin elements in adult specimens. *p*-values determined by Mann Whitney tests.

Alizarin red staining of the mineralized skeleton of *pax9^el622^*mutants and controls revealed that the expaned fin is fully supported by endoskeletal radials and exoskeletal rays (Fig. 2B, S2B’-E’). The median total number of endoskeletal elements plus fin rays across the anal and caudal fins was 77 in controls and 104 in mutants (*p*<0.0001, Mann-Whitney test; Fig. 2D). Considering external fin rays alone, the median was 42 in controls and 58 in mutants (*p*<0.0001; Fig. 2E). Individuals with a fully fused fin trended towards having more rays (median = 61) than those with a discontinuous fin (median = 53, but *p*>0.05; Fig. S2N). The short procurrent fin rays that demarcate the ventral border of the caudal fin were missing or reduced to one, even in specimens with a discontinuous ectopic fin (Fig. 2B’-C’, S2C’-E’,O). The caudal hypural elements ventral to the diastema were variably fused and misshapen (Fig. 2B’-C’, S2O). Most of the ectopic endoskeletal elements were not directly attached to vertebrae. Dorsal procurrent rays and hypural elements above the diastema looked relatively normal. The tips of both standard and ectopic fin rays are branched in the typical manner (Fig. 2B’). Quantification of dorsal, pectoral, and pelvic fin ray number also revealed a minor increase specifically in the dorsal fin, with a range of 9-10 rays in controls and 9-12 in mutants (*p*=0.0023, Fig. 2F-H). Some specimens were kinked at the vertebral terminus, but vertebrae were otherwise not dysmorphic, consistent with the *Pax9* mouse model in which vertebral phenotypes only emerge when *Pax1* gene dosage is also reduced (Peters et al., 1999).

### Normal upstream axial patterning in pax9 mutants

Two recent reports in zebrafish and medaka found that select combinatorial Hox gene mutations result in similarly dramatic elongation of median fins along the anterior-posterior axis or their complete absence, (Adachi et al., 2024; Cumplido et al., 2024). These fin phenotypes were linked to changes in axial patterning that also manifest as shifted vertebral number and identity. Of particular interest, *hoxc12a;c13a;c12b;c13b* zebrafish mutants develop a posteriorly extended anal fin (though it was not fused with the caudal fin (Adachi et al., 2024)), while *hoxb13a;c13a* mutants lack the caudal fin entirely (Cumplido et al., 2024). Given its timing of expression, we hypothesized that *pax9* works downstream of the Hox code. However, to exclude shifts in axial identity, we assayed two representative caudal Hox genes, *hoxd12* and *hoxc13a*, in *pax9* mutants by HCR at 24 hpf (Fig. S3A-C). The length of each gene’s domain in the paraxial mesoderm was indistinguishable between controls and mutants (Fig. S3D-E). Likewise, we detect no differences in axial segment number, determined by counting somites at 30 hpf and tail myomeres at 4 dpf (Fig. S3F-G). Expansion of the fin along the anterior-posterior axis in *pax9* mutants is thus unlikely to be a result of disrupted Hox coding.

### The ectopic fin emerges after formation of mispatterned caudal and anal fins

Formation of the ectopic ventral fin was surprising in both degree and location: first, because *Pax9* mutant mice develop only a single extra digit (one phalange plus expanded metatarsal/carpal cartilage)(Peters et al., 1998); second, because in mutants with a discontinuous phenotype, the anal fin was *posteriorly* elongated (Fig. S2) despite expressing *pax9* only at the anterior part of the bud (Fig. 1G). Seeking clues from the ontogeny of the phenotype, we collected mutants and sibling controls at larval through juvenile stages (6-30 dpf) and stained them with Alcian blue (cartilage) and Alizarin red (bone) (Fig. 3A-H). Cartilaginous endoskeletal elements of the caudal fin normally begin appearing between 3.7 and 4.3 mm SL (>8 dpf) (Bird & Mabee, 2003; Desvignes et al., 2022). No precocious development was apparent in mutants (Fig. 3A’). However, once they did appear, elements anterior to the future diastema were always truncated, misdirected, or fused with each other (Fig. 3B’-D’). Posterior elements looked largely normal. This suggests that the anterior domain of the mutant caudal fin bud is already abnormal before chondrocytes begin to differentiate. Indeed, imaging of *fli1:EGFP*+ bud mesenchyme revealed that the mutant bud is significantly larger in area than controls by 5 dpf (*p*<0.0001, Mann Whitney test; Fig. 3J-K), though no overt differences were detected at an early stage of bud development, 60 hpf (Fig. 3I).

**Fig. 3.**
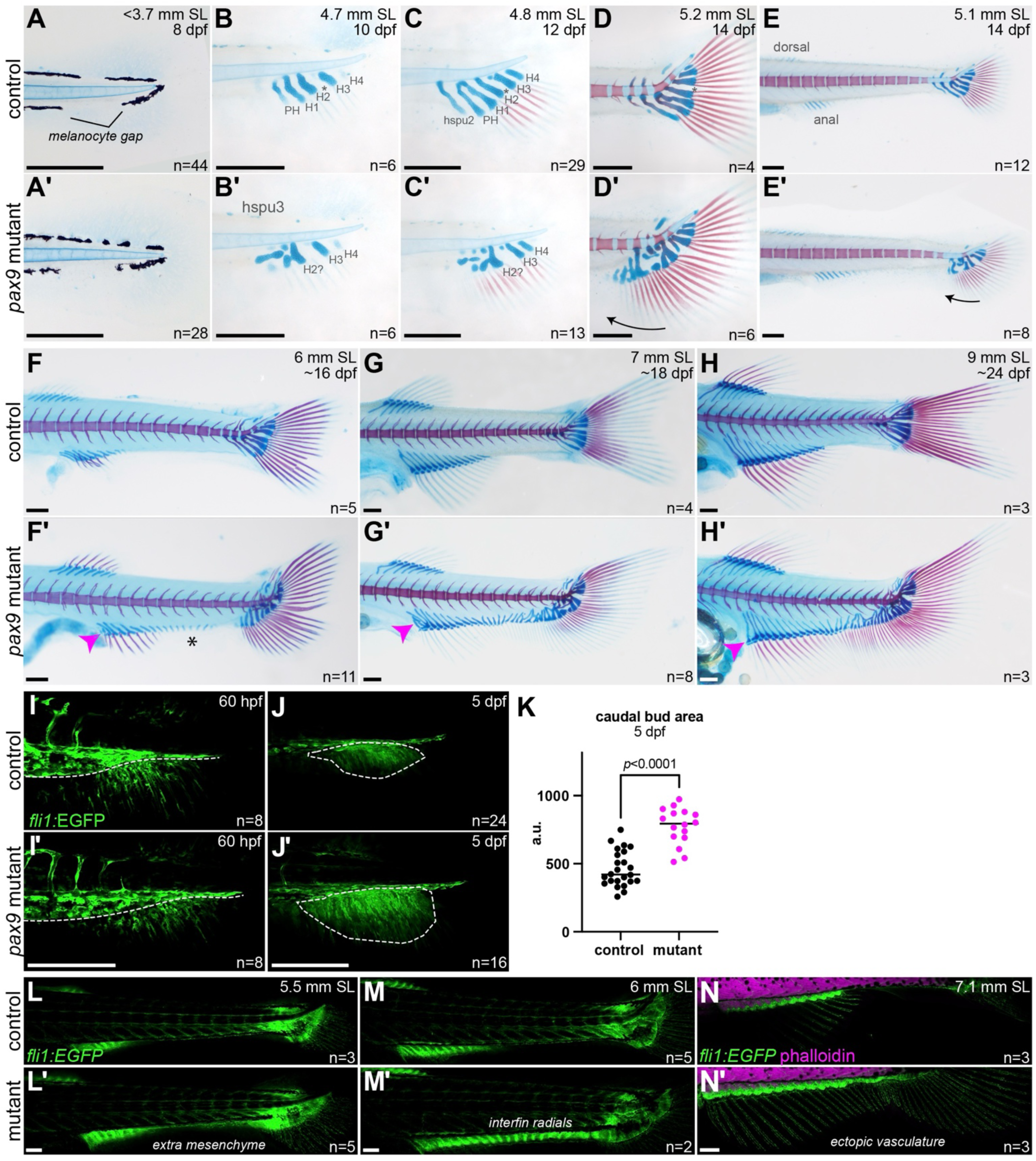
Ectopic fin fills in after formation of mispatterned caudal and anal fins. **A-H**, Alcian blue and Alizarin red staining of the developing median fins in larval and juvenile control and *pax9^el622^* mutant fish. Note the lack of precocious cartilage differentiation (A’), the variably branched and fused anterior hypural elements anterior to the diastema (asterisk) (labeled in B-C’ as in Fig. 1 legend), progressive anterior expansion of the caudal fin skeleton (marked with arrows in D-E’), and belated appearance of ectopic radials (F’) then rays (G’) in the interfin region (asterisk in F’). The adult pattern becomes evident between 7-9 mm SL (G’-H’). The youngest specimens shown in A-A’ (8 dpf) were left unbleached so to display the melanocyte gap. Magenta arrowheads mark mispatterned radials in the anterior anal fin. **I**, *fli1*:EGFP+ mesenchyme has begun to accumulate below the caudal vasculature at the site of the caudal bud by 60 hpf in controls and mutants (border indicated by dashed line). **J**, By 5 dpf, the EGFP+ bud domain is more well-defined and enlarged in mutants (J’). **K**, Quantification of caudal bud area at 5 dpf. a.u., arbitrary units. Significance determined by Mann-Whitney U test. **L-M**, Ectopic *fli1:*EGFP+ mesenchyme appears in the interfin region by 5.5 mm SL (L’), preceding formation of EGFP+ radials ∼6 mm SL (M’). **N**, The ectopic fin is fully vascularized by *fli1:*EGFP+ endothelial cells. Phalloidin staining marks muscle. Confocal images are maximum intensity projections. Scale bars = 250 μm.

Mutants had 6-9 extra caudal fin rays by ∼5 mm SL (∼14 dpf; approx. 26 versus 18 rays in controls, Fig. 3D-E), reminiscent of mammalian polydactyly. Anal fin radials emerged around 5 mm SL in both controls and mutants (Fig. 3E). Shortly thereafter, around 6 mm SL (∼16-17 dpf), ectopic anal fin-like radial elements not attached to the vertebral skeleton began to populate the interfin gap (Fig. 3F’). Confocal imaging of mutants carrying *fli1:EGFP* revealed that these ectopic elements emerge from an ectopic population of EGFP+ fin bud mesenchyme in the ventral fin fold (Fig. 3L-M). By 7-8 mm SL (17-20 dpf), abnormally patterned cartilaginous elements populated the entire interfin and lower hypural regions in mutants (Fig. 3G’). The ectopic fin is also fully vascularized by *fli1:*EGFP+ endothelial cells (Fig. 3N). The anterior-most radials of the anal but not dorsal fin were also branched or fused in most mutants imaged at or after 6 mm SL (Fig. 3F’-H’). By 9-10 mm SL (21-30 dpf), all external fin rays had ossified (Fig. 3H). At this stage, the adult mutant phenotype is fully realized, with continuous skeletal elements present along the ventral midline (Fig. 3H’).

### Missing anterior boundary and posterior character of expanded fins

We hypothesized that the dysmorphic elements in the anterior anal and caudal fins are a sign of disrupted anterior identity within the buds. To test this, we crossed in the *alx4a:DsRed* BAC transgene (Nachtrab et al., 2013) along with osteoblast marker *sp7:EGFP* (DeLaurier et al., 2010) to label the fin rays (Fig. 4A-C). At 7-9 mm SL, mutants with a fully fused ventral fin lacked the bright *alx4a*:DsRed+ stripe marking the ventral (originally anterior) caudal fin edge in controls (Fig. 4A’). *alx4a*:DsRed+ domains in the anterior anal fin and dorsal (originally posterior) caudal fin were present but weakened, while those in the dorsal and pectoral fins appeared less affected. Earlier, at 5.5 mm SL, all mutants showed supernumerary elements in the caudal fin, and most but not all completely lacked *alx4a*:DsRed activity at the anterior boundary, with relatively normal levels in the other domains (Fig. 4B-C, S2P).

**Fig. 4.**
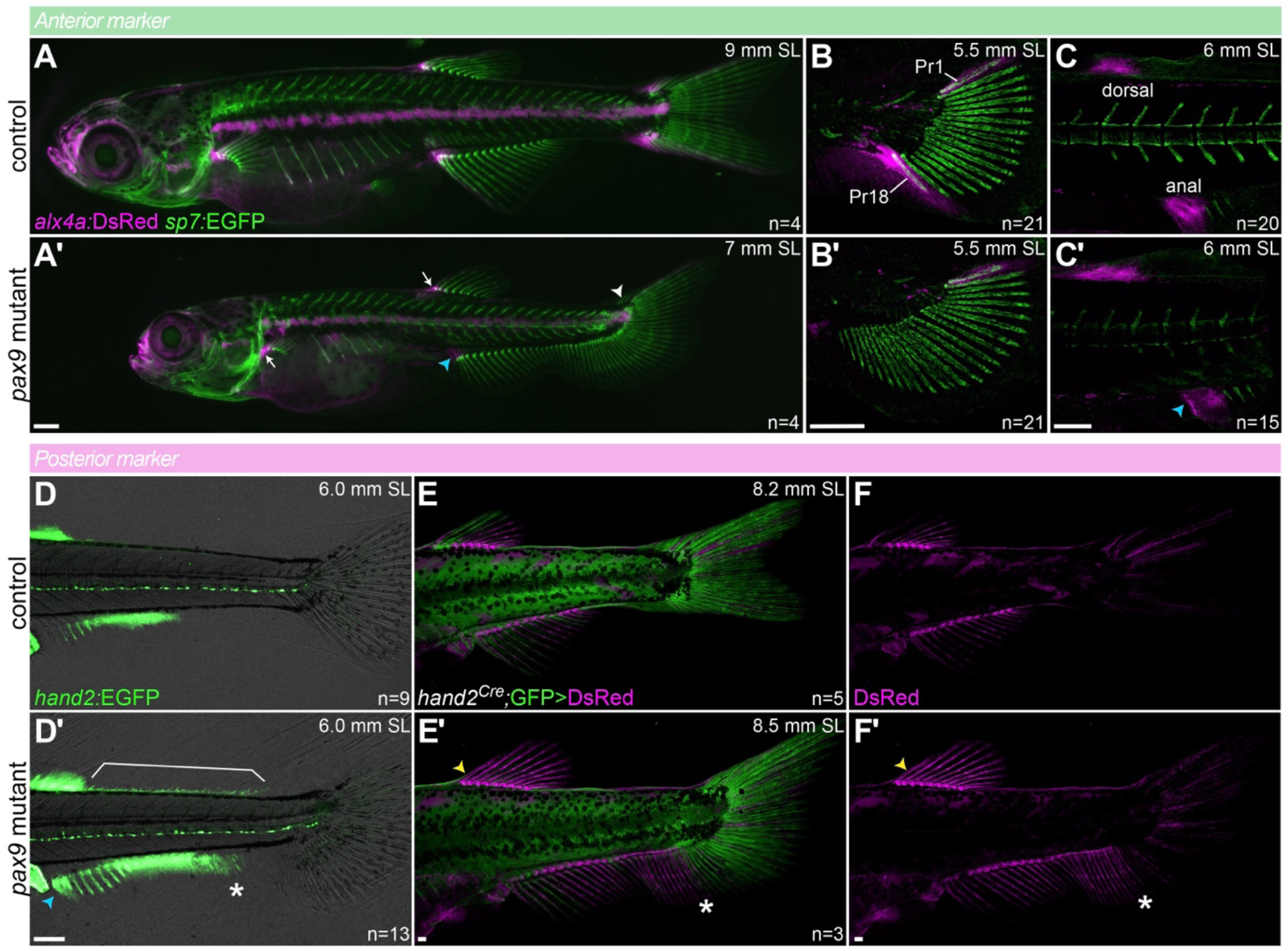
Missing anterior boundary and posterior character of expanded anal fin. **A**, Relative to sibling controls, juvenile *alx4a:DsRed*; *sp7:EGFP pax9^el622^* mutant fish have reduced DsRed signal in the anterior anal fin (blue arrowhead) and upper caudal fin (white arrowhead), no signal within the fused caudal/ectopic fin, and fairly normal levels in the dorsal and pectoral fins (white arrows). **B-C**, Loss of *alx4a:*DsRed signal around the EGFP+ anterior-most principal fin ray 18 (Pr18) is evident as early as 5.5 mm SL. Fluorescence is still evident around Pr1 in the unaffected posterior region. DsRed is also reduced in the anal but not dorsal fins at 6 mm SL (C’). **D**, Relative to sibling controls, juvenile *hand2:EGFP* mutant fish show ectopic EGFP signal in the ectopic fin (asterisk in D’), the anterior anal fin (blue arrowhead) and dorsal body margin (bracket). Transmitted light channel is shown for context. **E-F**, Lineage tracing of *hand2*-expressing cells with the *ubi:Switch* GFP>DsRed reporter reveals labeling of most of the ectopic fin skeleton (asterisk in E’-F’) and an anterior shift in the dorsal fin boundary (yellow arrowheads). F-F’ show Cre-converted cells only. Images are maximum intensity projections. Scale bars = 200 μm.

To test whether the ‘posterior’ program is erroneously activated when anterior identity is impaired, we crossed in reporters for *hand2* (Fig. 4D-F), a conserved posterior marker (Charite et al., 2000; Yelon et al., 2000). As reported (Nachtrab et al., 2013), *hand2:*EGFP activity is absent in the wild-type caudal fin (Fig. 4D). This remained the case in mutants (n=13/13; Fig. 4D’). However, the central section of the ectopic fin was *hand2:*EGFP+ (n=13/13), as was the anterior-most domain of the anal but not dorsal fin (n=10/13 anal, 0/5 dorsal; Fig. 4D’). Disrupted molecular identity of the anterior anal fin agrees with its mildly aberrant pigmentation pattern (Fig. S2C-E,G,I). Ectopic *hand2:*EGFP+ cells were also observed lining the dorsal body margin between dorsal and caudal fins in the seven younger mutants (<6.0 mm SL; Fig. 4D’). A knock-in *hand2^Cre^*(Ming et al., 2026) crossed with the *ubi:LOXP-GFP-LOXP-DsRed* (*ubi:Switch*) reporter (which converts from GFP to DsRed) (Mosimann et al., 2011) produced similar patterns: no DsRed+ cells in the caudal fin, strong DsRed signal in the posterior anal/ectopic fin up to the point of fusion with the expanded caudal fin, and expansion of DsRed into the anterior of both anal and dorsal fins (Fig. 4E-F). Putting these reporter patterns together with the skeletal phenotypes, *pax9* is essential for enforcing the anterior boundary of the caudal fin and for supporting anterior identity in the caudal, anal, and – to a lesser degree – dorsal fins. In the paired fins, by contrast, *pax9* is expressed but apparently dispensable.

### The ectopic fin derives from the paraxial mesoderm

We next considered the cellular origins of the ectopic fin. One possibility is spreading of caudal fin bud mesenchyme due to loss of the anterior boundary, supported by early overgrowth of the bud (Fig. 3J-K) and caudal fin (Fig. 3C’-E’, 4B’) prior to emergence of the ectopic fin. However, this is not consistent with the posterior elongation of the anal fin in the discontinuous mutants (Fig. S2), nor with the activation of the *hand2:GFP* reporter in most of the ectopic fin (Fig. 4D’). Another option is abnormally-behaving trunk neural crest cells: a recent study found that misexpression of the oncogenic EWSR1::FLI1 fusion protein reprogrammed neural crest cells to a mesoderm-like state, resulting in the production of small ectopic fins throughout the body (Vasileva et al., 2025). To determine the neural crest contribution to the ectopic ventral fin of *pax9* mutants, we performed lineage tracing with *SOX10:Cre* in combination with the *actb2:LOXP-BFP-LOXP-DsRed* Cre reporter (which converts from BFP to DsRed) (Kague et al., 2012; Kobayashi et al., 2014). Scattered lineage-labeled DsRed+ cells were present in the ectopic fin but resembled glia or pericytes, not skeletal elements or mesenchyme (Fig. 5A-B). These data indicate that our mutant’s ectopic fin does not originate from neural crest cells.

**Fig. 5.**
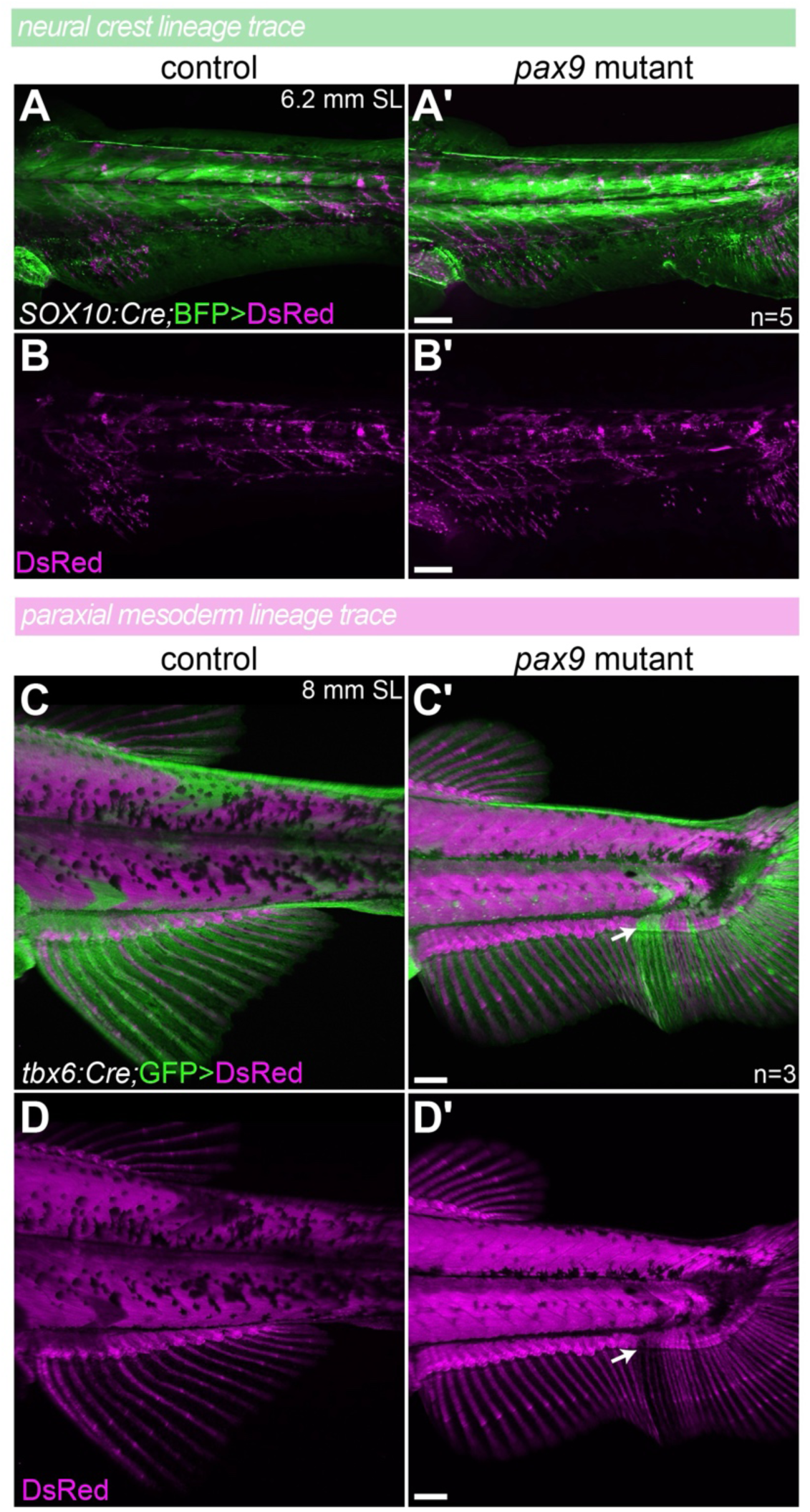
The ectopic fin derives from the paraxial mesoderm. **A-B**, Lineage tracing with neural crest-specific *SOX10:Cre* and the *actb2:BFP>DsRed* reporter reveals only a few converted cells in the ectopic fin, likely associated with vasculature. B-B’ show Cre-converted cells only. **C-D**, Lineage tracing of paraxial mesoderm with *tbx6:Cre* and the *ubi:Switch GFP>DsRed* reporter confirms that the ectopic fin skeleton derives from the somites. Note the mosaically unconverted EGFP+ somite in the mutant directly adjacent to a swath of GFP+ fin (arrow). D-D’ show Cre-converted cells only. Images are maximum intensity projections. Scale bars = 200 μm.

As a third possible source, we considered ectopic contribution from somites that do not normally make fin. Cells from the sclerotomal compartment of the somites form the endogenous median fin skeleton, as definitively demonstrated by lineage tracing with *tbx6:Cre* (paraxial mesoderm) and *nkx3-1:Gal4;UAS:CreERT2* (sclerotome) as well as somite transplants (Lee et al., 2013a; Lee et al., 2013b; Shimada et al., 2013; Bailon-Zambrano et al., 2024). Using *tbx6:Cre* with the *ubi:Switch* Cre reporter (Fig. 5C-D), we observed broad DsRed+ lineage-labeling of the ectopic fin skeleton, supporting that it too derives from the paraxial mesoderm. Of even greater interest, however, were the sporadic mosaically unconverted segments, in which an EGFP+ myomere connected to an EGFP+ section of the ectopic fin (Fig. 5C’-D’, arrows). These indicate that that the skeletal mesenchyme in the interfin region derives from directly adjoining somites, not solely from spreading of the caudal (or anal) fin bud.

### Shift in a sclerotome fate decision may explain formation of ectopic fin

To obtain a comprehensive view of the cell type composition and gene expression changes altered by *pax9* loss of function, we performed scRNA-seq analysis of pooled 3 dpf mutant and closely related wild-type control tails (Fig. 6A). After filtering, there were 12,901 mutant and 11,764 wild-type cells, divided by marker gene analysis into periderm (2 clusters), epidermis (2), spinal/glial cells (3), mesenchyme (2), lateral line (2), hematopoietic cells (1), fin fold fibroblasts (1), notochord (1), and muscle (1) (Fig. 6B, Table S1). No clusters were exclusively comprised of one genotype or the other (Fig. S4A-B), and no cell types segregated by proliferation status except the mesenchyme (Fig. S4C). Minimal differentially expressed genes were detected by Wilcoxon rank-sum testing (22 upregulated, 38 downregulated; log2 fold-change >0.585 or <-0.585; adjusted *p*<0.05; Fig. S4D; Table S2). Twelve of the upregulated and 9 of the downregulated genes had mesenchyme-enriched expression, but the changes in expression level were mild.

**Fig. 6.**
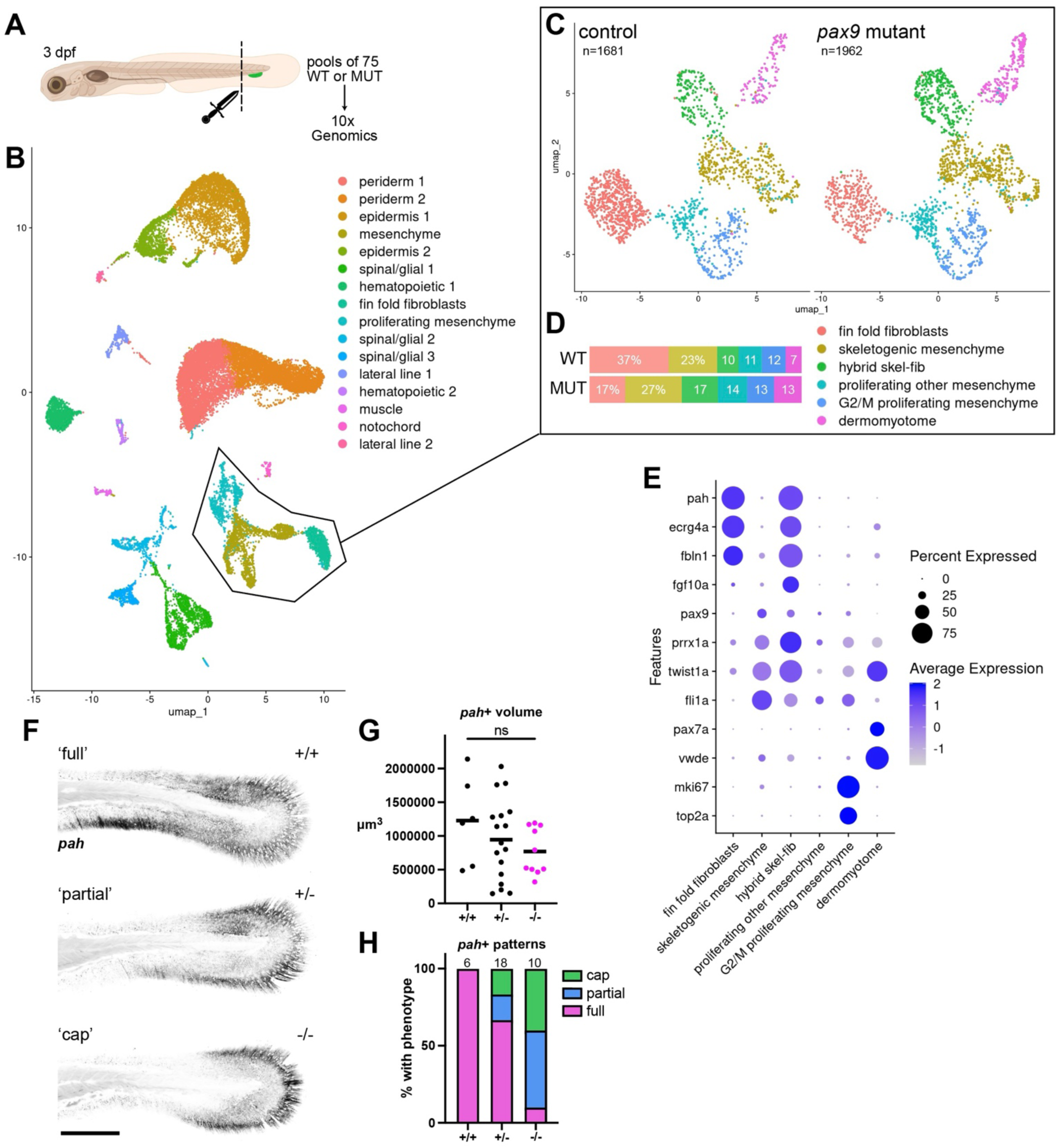
Shift in a sclerotome fate decision may explain formation of the ectopic fin. **A**, scRNAseq sample preparation. Related all-mutant or all-wild-type clutches were generated through selective breeding, and tails were removed from 3 dpf larvae with a micro-knife and pooled (75/sample) prior to creating cell suspensions for 10x Genomics analysis. **B**, UMAP of pooled wild-type and mutant cells divided into 16 clusters. Cell types were ascertained by marker gene analysis. **C**, Mesenchyme and fin fold fibroblast clusters were subsetted and reanalyzed, then projected into UMAPs separately by genotype. **D**, Proportions of each sample that fell into each of the six new clusters. **E**, DotPlot for selected marker genes defining each of the six clusters. **F**, Examples of the three patterns of *pah+* fin fold fibroblasts in the tail revealed by HCR staining. Images are maximum intensity projections. **G**, Quantitation of *pah*+ volume by genotype. Significance determined by one-way ANOVA followed by Dunn’s multiple comparisons test. **H**, Proportion of each genotype showing the indicated pattern.

We then narrowed our focus to the mesenchymal cells and fibroblasts, which yielded six new clusters upon subclustering: fin fold fibroblasts (*fbln1*+, *pah*+, *ecrg4a*+), skeletal mesenchyme (*fli1*+, *twist1a*+, *prrx1a*+, *pax9*+, *pdgfra*+), dermomyotome (*pax7a*+, *vwde*+), two proliferating mesenchyme clusters, and a distinct *fgf10a*+ hybrid cluster expressing markers of both skeletal mesenchyme and fin fold fibroblasts (Fig. 6C,E, S5A-C, Table S3). The hybrid cluster may be a remnant of the *fgf10a*+ tail bud (Fig. S6A-B) (Thisse & Thisse, 2004; Lalonde & Akimenko, 2018). Interestingly, fin fold fibroblasts made up a much greater proportion of the wild-type than the mutant sample (37% vs. 17%) (Fig. 6D). All other clusters were more highly represented in the mutant, particularly skeletal mesenchyme, consistent with the enlarging caudal fin bud and later production of ectopic fin skeleton in the interfin region. Within the skeletogenic mesenchyme cluster, we distinguished a subpopulation of likely vertebral precursors (*nkx3-2*+, *pax1a*+, *cdh11+*, *col9a3*+) (Rajan et al., 2023) from anterior (*pax9+*, *alx4a+*) and posterior (*grem1b*+) fin bud (Fig. S5D).

Differential expression analysis of the mesenchymal cluster with standard Wilcoxon rank-sum tests identified only 13 genes downregulated in mutants relative to wild-type, versus 146 upregulated (Fig. S7A-B, Table S4). We suspected this asymmetry was a technical artifact caused by higher detection rates in mutant cells (mean_features = 1176 in wild-type vs. 1359 in mutant; mean_counts = 3351 vs. 3709, respectively). We thus repeated the analysis using a separate linear regression model compatible with covariate adjustment. Including nFeature_RNA and nCount_RNA in the model reduced but did not eliminate the imbalance between up- and downregulated genes (dropping from 16 to 13 downregulated and 167 to 50 upregulated; Fig. S7B). This indicates that differences in gene detection sensitivity did influence the original result. However, across all lists, the rare downregulated genes were all nearly exclusively expressed in fin fold fibroblasts and the hybrid population, while upregulated genes (e.g. *ccn1*, *jund*, and *pax9* itself) were broadly distributed across the other cell types (Fig. S7C-D). We validated the predicted *pax9* upregulation by in situ at 3 dpf, noting an anterior expansion of the *pax9*+ caudal fin domain inside the fin fold (Fig. S7E), in line with the expanding caudal fin bud (Fig. 3J-K).

The apparent depletion of the fin fold fibroblast population and suppression of fin fold fibroblast genes in mutants were intriguing because, like the median fin skeleton, most fibroblasts of the median fin fold derive from the sclerotome (Ma et al., 2023). However, they move out to the periphery much earlier, between 26 and 30 hpf (Lee et al., 2013a). Both the sclerotome itself and the migrating fibroblasts initially express *pax9* (Fig. 1H-K) (Ma et al., 2018; Sur et al., 2023). To validate the predicted reduction of fin fold fibroblasts in mutants, we performed HCR for a specific marker, *phenylalanine hydroxylase* (*pah*) (previously characterized in dermal fibroblasts (Farmer et al., 2024)), and measured the volume of fin fold covered by *pah*+ cells in tails of 3 dpf larvae (Fig. 6F-G). Differences in *pah*+ volume between genotypes were not significant (Fig. 6G), due in part to variable staining intensity. High cell density and the surrounding epithelial layers impeded reliable cell counting at this stage. However, three qualitative *pah* expression patterns emerged: ‘full’, ‘partial’, and ‘cap’, in order of decreasing ventral area covered (Fig. 6F). The cap corresponds to the part of the tail that does not pass through a *pax9/nkx3-1*+ sclerotome state (see Fig. S1B) (Ma et al., 2023). The ‘full’ pattern was observed in all wild-type and most but not all heterozygotes (Fig. 6H). Mutants primarily presented the cap and partial patterns, with overtly reduced *pah* staining – i.e., fewer fin fold fibroblasts – in the ventral fin fold where the ectopic fin later develops (Fig. 6H). The mutant fin fold itself was not morphologically abnormal (Fig. S6C). These findings suggest that the later increases in fin bud mesenchyme within the caudal and interfin regions (Fig. 3J,L-M) may come at the expense of the fin fold fibroblast population.

To determine whether the sclerotome itself was overtly abnormal in *pax9* mutants, we performed HCR for the pan-sclerotome marker, *nkx3-1*, at 24 hpf, prior to fin fold fibroblast emergence. *nkx3-1* activates earlier than *pax9* and is not thought to be a target (Ma et al., 2018). The volume of *nkx3-1*+ sclerotome was measured in the posterior trunk, starting 5 somites anterior to the end of the yolk extension (Fig. S8A-B). No differences in volume (p=0.23; Fig. S8C) or qualitative appearance of the sclerotome were noted in mutants at this stage. This suggests that the problem is occurring somewhat later as sclerotomal derivatives begin to differentiate, manifesting as an altered balance between skeletal mesenchyme and fin fold fibroblasts by 3 dpf in our scRNAseq data. The key downstream targets through which Pax9 works in the sclerotome may thus no longer be detectable or dysregulated in our scRNAseq dataset. However, we reasoned that its downstream targets in the anterior caudal fin bud could be identifiable.

We therefore re-ran the differential expression analysis specifically on *pax9*+ cells and found 10 genes significantly downregulated vs. 3 up between mutants and controls (Wilcoxon rank-sum test, Fig. S9A, Table S5). *pax9* itself was still called as upregulated, along with *bmpr1ba* and *angptl2a* (Fig. S9A,C,E). *tbx3a*, a homolog of a mammalian limb patterning gene (Emechebe et al., 2016), was sharply downregulated, as was a Wnt inhibitor, *sfrp1a* (Fig. S9A-B,D). Both genes are enriched in the putative ‘anterior fin bud’ subcluster that co-expresses *pax9* and *alx4a* (compare Fig. S9D to S5D and S7D). We next scanned a published PAX9 ChIP-seq dataset from murine vertebral cells (Sivakamasundari et al., 2017) for evidence of binding near the mouse homologs of these dysregulated genes. Nearest-gene analysis of BED data using GREAT (McLean et al., 2010; Tanigawa et al., 2022) identified 24 PAX9 ChIP-seq peaks around the *Pax9* locus (supporting that it auto-represses its own transcription), 7 near *Bmpr1b*, 3 near *Angptl2*, 17 near *Tbx3*, and none near *Sfrp1* (Table S5). Future studies will investigate the activity and conservation of these putative enhancer sequences as well as the involvement of the target genes themselves in median fin patterning.

## DISCUSSION

Our findings lead us to propose that Pax9 plays an unusual dual role in median fin development (Fig. 7): As anticipated, it promotes anterior identity in all median fins (albeit weakly in the dorsal fin) and enforces the anterior boundary specifically in the caudal fin. Loss of this function disorganizes the anterior endoskeleton of both anal and caudal fins, eliminates the anterior procurrent rays of the caudal fin, and increases caudal fin ray number. Intriguingly, the *pax9*+ anterior boundary of the caudal fin forms directly beneath the most posterior two somites to develop a bona fide *pax9*+ sclerotome (see Figs. 1L, S1B, 7, left). It is conceivable that these two events are interdependent, i.e. that those posterior-most *pax9*+ sclerotomal cells migrate ventrally to the base of the fin fold and concurrently establish the anterior boundary and identity of the nascent caudal fin bud. Though better lineage-tracing tools are needed to rigorously test this hypothesis, it offers an appealing new mechanism for ZPA-independent anterior-posterior patterning of the caudal fin and aligns with the primacy of Pax9 in this particular appendage.

**Fig. 7.**
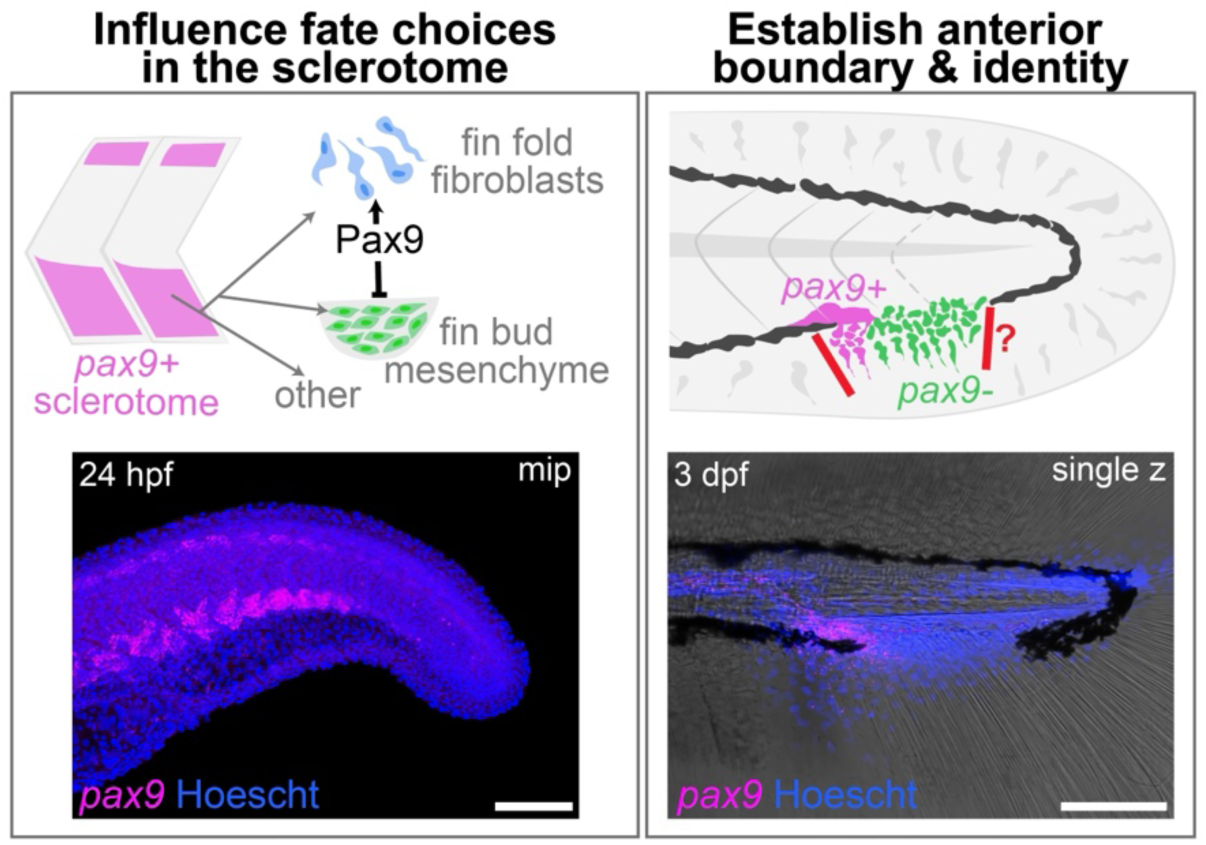
Summary of Pax9’s dual role in median fin development. Left, Pax9 acts early within the sclerotome to influence a putative fate choice between fin fold fibroblasts and fin mesenchyme, thus capping the fin progenitor population. Right, Pax9 also acts later in the caudal fin bud to activate the anterior identity program and enforce the anterior boundary.

The second part of Pax9’s dual role actually occurs earlier, when it acts within the sclerotome to influence the differentiation of the mesenchymal cells that will populate the fin fold. This includes those cells that emigrate early and become fin fold fibroblasts, as well as those that accumulate later to form the fin buds (Fig. 7, right). In *pax9* mutants, the balance of fate choices within the sclerotome shifts away from fin fold fibroblasts, and normally ‘finless’ somites ectopically deploy fin bud mesenchyme to the ventral midline to seed the ectopic fin. There is evidence that this occurs dorsally too, albeit more subtly: many mutants develop 1-2 extra dorsal fin rays (Fig. 2F), and ectopic *hand2:*EGFP+ cells are present along the dorsal midline (Fig. 4D’). However, we never detected ectopic *fli1:*EGFP activity there, suggesting that the ectopic *hand2:*EGFP+ cells have not advanced to bona fide fin bud mesenchyme. This milder dorsal phenotype might be explained by the smaller size of the dorsal sclerotomal compartment (Ma et al., 2018) or requirements for additional, unidentified gene(s) to induce fin bud mesenchyme in this domain. Based on mouse data (Peters et al., 1999), we expect that zebrafish Pax9 is also required within an additional sclerotomal subcompartment – vertebral precursors – but here it likely works redundantly with its homolog Pax1a, which is specifically enriched in this subpopulation and excluded from the fin (Fig. S5D) (Ma et al., 2018).

These dual roles for Pax9 in median fin development might stem from the same molecular function in sclerotome and in the anterior fin bud: restraining the production of fin bud mesenchyme. Our data suggest that, in the absence of Pax9, early sclerotomal progenitors produce an excess of fin bud mesenchyme at the expense of fin fold fibroblasts. This impacts every segment and eventually leads to formation of the ectopic fin. Later, the mutant caudal fin bud overgrows in the anterior direction where *pax9* is normally expressed, in an analogous overproduction of fin bud mesenchyme. Either of these actions could account for the elevated proportion of mesenchyme we detect in the scRNAseq data. This interpretation is also supported by the *Pax9* mouse mutant phenotype, in which excess mesenchyme in the anterior region precedes formation of a preaxial digit in each limb (Peters et al., 1998). Why we detect no role for *pax9* in the paired fins is unclear, but may be explained by a later onset or lower level of expression in pectoral fin buds.

Which somites normally contribute mesenchyme to the median fins and which are ‘finless’ has been mapped in medaka by somite transplant experiments (Shimada et al., 2013). Mutant studies have further revealed that this blueprint is imposed by combinatorial Hox gene function (Adachi et al., 2024; Cumplido et al., 2024). However, how that blueprint is interpreted to initiate budding of mesenchyme only at specific axial positions has remained an open question. Our results indicate that Pax9’s activity within the sclerotome is part of the answer, yet we still must reconcile the fact that *pax9* is expressed in the sclerotome of every somite that compartmentalizes, not just those that do or do not make fin. One possibility is that Pax9’s interaction with target gene enhancers is altered by the presence of different Hox factors at different axial positions. For instance, Pax9 could repress a skeletal gene enhancer except in the presence of certain Hox factors. In this case, the gene would be de-repressed everywhere when Pax9 protein is missing. Such interactions between Pax and Hox factors have been previously demonstrated, e.g. PAX3 and HOXB6 in control of a *Myf5* enhancer during rib development (Guerreiro et al., 2013) and PAX2 and HOX11 on a *Six2* enhancer in the kidney (Gong et al., 2007).

We cannot find a natural example of a fish mimicking the fused caudal and anal fins of the zebrafish *pax9* mutant, indicating that this is a disadvantageous form. Lungfish, eels, and tonguefish develop a continuous median fin that incorporates the dorsal, caudal, and anal domains, but the structure is highly simplified and lacks regional character. Other comparisons are likewise imperfect: the anal fins of Betta fish are elongated posteriorly but remain separate from the caudal fin. Knifefish also have an elongated anal fin, but the caudal fin has been lost. In general, having the caudal fin attach to the rest of the body through a slender, finless stem is thought to reduce viscous drag during sustained swimming and thus optimize propulsion (Webb, 1984). Though our mutants do not show overt swimming deficits in the laboratory, reduced propulsion and/or recurrent damage to the tissue from shear forces could select against this form in nature.

Our finding that Pax9 regulates production of the sclerotome-derived mesenchymal cells that populate the median fin fold and median fin buds in zebrafish draws attention to how poorly we understand the evolutionary loss of these structures in our own lineage, despite the preservation of sclerotomal Pax9 activity. Early tetrapods lost their median fins in the late Devonian period as they adapted their locomotory strategies to the terrestrial environment. Speculation regarding the genetic and developmental changes leading to this loss of fins is sparse (though see (Cumplido et al., 2024)). We also have yet to determine what confers the sclerotomal cells of fish and chondrichthyans with the ability to move to the midline and initiate median fin bud formation. Though we have detected apparent contiguity between the sclerotome of the most caudal compartmentalized somites and the caudal fin bud, the dorsal and anal fin buds form much later in development. Where their sclerotome-derived progenitors reside in the interim and how they are summoned to initiate bud development remain open questions.

## MATERIALS & METHODS

All animal procedures were approved by the Institutional Animal Care and Use Committees of Cincinnati Children’s Hospital Medical Center (Nos. 2021-0048 and 2024-0108), the University of Colorado (No. 00188), Purdue University (No. 1210000750), and the University of Calgary (Nos. AC13-0236 and AC21-0102).

### Zebrafish lines and genotyping

*Danio rerio* (zebrafish) embryos were incubated in embryo medium at 28.5°C up to 5-6 dpf (Westerfield, 2007) and staged as described (Kimmel et al., 1995). Larvae and juvenile fish were maintained on different standard diets at the different institutions, including rotifers, paramecia, and Gemma (Skretting), with brine shrimp introduced at approximately two weeks post fertilization. Published lines used in this study include wild-type stocks Tübingen (ZDB-GENO-990623-3) (Haffter et al., 1996), AB (ZDB-GENO-960809-7) (Chakrabarti et al., 1983), and TL (Tüpfel long-fin; ZDB-GENO-990623-2); mutant alleles *pax9^el622^*, *pax9^ci3038^* (Paudel et al., 2025), and the transparent stock *absolute* (*ednrb1a^b140^; mitfa^b692^*) (ZIRC ZL1689); as well as transgenics *Tg(fli1:EGFP)^y1^*(Lawson & Weinstein, 2002), *Tg(alx4a:DsRed2)^pd52^*(Nachtrab et al., 2013), *Tg(hand2:EGFP)^pd24^* (Kikuchi et al., 2011), *Tg(sp7:EGFP)^b1212^* (DeLaurier et al., 2010), *Tg(hand2-2A-cre, gcry1:mRFP*)*^is65^* (aka *hand2:Cre*) (Ming et al., 2026), *Tg(Mmu.Sox10-Mmu.Fos:Cre)* (aka *SOX10*:*Cre*) (Kague et al., 2012), *Tg(actb2:LOXP-BFP-LOXP-DsRed)^sd27^*(Kobayashi et al., 2014), *Tg(tbx6:Cre)^sq6^* (Lee et al., 2013b), and *Tg(−3.5ubb:LOXP-EGFP-LOXP-mCherry)^cz1701^* (aka *ubi:Switch*) (Mosimann et al., 2011). Genotyping of established *pax9* mutant lines was performed as described (Paudel et al., 2025). We primarily used the *pax9^el622^*line for experiments after confirming that the primary median fin phenotype was also present in the *pax9^ci3038^*line. Both wild-type and heterozygous siblings were used as controls.

### Generation of new zebrafish lines

One new *pax9* CRISPR-Cas9 knockout line, *pu116*, was generated on the TAB background (AB/Tübingen, RRID: ZIRC_ZL1) following a previously described approach using two gRNAs targeting exon 2 (Dr.pax9-CR1: 5’-CCAACTCTACTATCCTGAGCCGG-3’; Dr.pax9-CR2: 5’-AATCGGCGGCAGTAAACCGAGGG-3’) (Kim & Zhang, 2020). The resulting *pax9^pu116^* allele carries a 77-bp deletion and a 43-bp insertion in exon 1. Two other new *pax9* mutant alleles, *ca122* and *ca123*, were recovered from a TL clutch injected with Cas9 and a gRNA targeting a more 3’ position in exon 2 (5’-GGTATGGTGGGTTGAGGTTG -3’). *pax9^ca122^* is a 20-bp insertion, and *pax9^ca123^* is a 7-bp deletion.

The *pax9* CRISPR-Cas9 knockin line, *pax9^hsp70l-mlanYFP^ ^pu115^*, was generated using a non-homologous end joining-based strategy modified from a previously published method (Kimura et al., 2014). A single gRNA targeting the 5’UTR of *pax9* (T7pax9-Ex1G1: GACTCGGAACAGGTCAGAAT) was injected into wild-type TAB zebrafish to mediate targeted integration of a plasmid containing the *hsp70l-mlanYFP* sequence. The full activation pattern of this allele, including expression in the paired fins, will be described elsewhere (Dong et al., in preparation).

Fish lines were verified by Sanger sequencing F1 adult fish.

### Skeletal staining

For developmental analyses, ontogenetic series were collected from large clutches and staged at key time points corresponding to significant milestones in median fin development. At each selected endpoint, zebrafish were fully anesthetized with 0.2% Tricaine (MS-222). Adults were then fixed in 4% paraformaldehyde (PFA) overnight at 4°C and juveniles in 2% PFA for 1.25-3 hours at room temperature. After fixation, samples were stored in PBS at 4°C until use. For larval and juvenile (8-24 dpf) stages, Alcian blue and Alizarin red staining was performed as previously described (Walker & Kimmel, 2007; Ullmann, 2011) with bleaching omitted at 8 dpf. Smaller samples were mounted in 50% glycerol and then imaged with a compound light microscope (Nikon Eclipse Ni-E), acquiring 5-6 optical z-sections per sample, which were subsequently combined into an extended-focus projection. Larger samples (≥5 mm SL, ≥14 dpf) were mounted in 50-75% glycerol and imaged using a stereo microscope (Zeiss Discovery V8). Standard length (SL) was recorded manually at time of imaging for samples over 15 dpf, measured from the tip of the snout to the end of the notochord or base of the caudal fin. Samples under 15 dpf were measured digitally with Nikon Elements software using the same measurement parameters.

As it was challenging to thoroughly clear large adult samples with trypsin and potassium hydroxide while preserving delicate fin rays, we adapted a published RAP (“rapid and nondestructive”) protocol (Sakata-Haga et al., 2018). Standard adult 4% PFA fixation was performed overnight at 4°C. After PBS washing, samples were immersed in 1% KOH, 5% Triton X-100 for 24-48 hours at 42°C with gentle rocking. This solution was exchanged after 24 hours. For further tissue clearing, samples were then directly immersed in a solution of 20% ethylene glycol, 1% KOH, 5% Triton X-100 for increased optical clarity and depigmentation. After incubation at 42°C for 12-48 hours, fish were briefly washed in PBS to remove scales. The samples were then submerged in 20% ethylene glycol, 1% KOH for 15 minutes, followed by Alizarin red staining (0.05% Alizarin, 1% KOH, 20% ethylene glycol) for 30 min-1 h at room temperature. Destaining of soft tissue in 1% KOH, 20% Tween was performed at 42°C for 4-12 h. Samples were stored in 50% glycerol until imaging, at which point they were mounted in 0.5% agarose, and brightfield images were captured with a stereomicroscope (Zeiss Discovery V8).

### Quantification and categorization of median fin elements

To systematically identify and score ectopic skeletal elements, a standardized scoring rubric was developed based on published descriptions of wild-type zebrafish skeletal morphology and executed in Fiji/ImageJ using the manual multipoint (spot count) feature. All median and paired fin skeletal elements were quantified using the same consistently applied process. Standard length (mm SL), developmental stage (dpf), and clutch were recorded for all samples to account for allometric effects. All counts presented herein are for adult specimens only. Ectopic elements were defined as those absent in wild-type fish, including ectopic proximal and distal anal radials, ectopic caudal hemal spines, and ectopic fin rays. In wild-type fish, the six vertebrae anterior to the posterior vertebral complex lack associated radials or hemal spines (Bird & Mabee, 2003). Accordingly, any skeletal elements affiliated with these normally finless vertebrae were classified as ectopic. Both procurrent and primary rays were included in the fin ray counts shown in Fig. 2. For blinding purposes, samples were genotyped after imaging and initial quantification, although the phenotype is unmistakable by 14 dpf. Degraded samples with distorted morphology or missing fin rays were excluded from quantification. As most datasets did not present normal distributions, genotypes were compared using non-parametric Mann-Whitney tests, with significance defined as *p*<0.05. Graphs were produced in Prism 11 (GraphPad).

### In situ hybridization

Colorimetric in situ hybridization was performed using a previously published probe for *pax9* (Paudel et al., 2022) following an all-age protocol that uses xylol and acetone for permeabilization (Vauti et al., 2020). The pigmentless *absolute* line was used for early-stage samples. *pax9^el622^*embryos were genotyped in advance due to staining substrates inhibiting the PCR reaction. Specimens were imaged on a stereomicroscope (Zeiss Discovery V8).

Hybridization chain reaction in situs were performed according to the manufacturer’s protocol (Molecular Instruments) using manufacturer-designed probes for *fgf10a, hoxc13a*, *hoxd12*, *nkx3-1*, *pah*, and *pax9*. Proteinase K treatment was skipped for embryos <24 hpf but applied at 30 μg/ml for 15 min at 2 dpf, 30 min at 3 dpf, and 50 min at 7-8 dpf. Probes were added at 10 μl/500 μl buffer except for *nkx3-1* (3 μl/500 μl) and *pax9* (2 μl/500 μl). Samples were imaged on a Nikon C2 confocal or a Leica DMi8 microscope equipped with an Andor Dragonfly 301 spinning disk then genotyped. Proteinase K was added to the lysis reaction at double the normal dose to facilitate genotyping post-HCR.

### Other fluorescent imaging

Alexa647-conjugated phalloidin staining was performed on fixed embryos as described (Barske et al., 2020), and samples were imaged on a Nikon C2 confocal microscope. Juvenile *alx4a:DsRed;sp7:EGFP* whole-body images were taken live under Tricaine anesthesia on a StereoDiscovery V8 fluorescent dissecting microscope (Zeiss) as separate channels later merged in Adobe Photoshop. All other fluorescent transgenic fish lines were imaged live under Tricaine anesthesia on a Nikon C2 or Andor Dragonfly confocal.

### Imaging data analysis

Nikon z-stacks were processed using Denoise AI, converted to maximum intensity projections, and exported as .tifs. Images were further processed in Adobe Photoshop, with all changes applied evenly across the entire image. The lengths of the *hoxc13a* and *hoxd12* domains in 24-hpf embryos were measured with the segmented line tool in ImageJ. Duplicate measurements were taken and averaged by an analyst blind to genotype. The area of the caudal fin bud was measured in ImageJ as the area occupied by *fli1*:EGFP+ cells ventral to the caudal vasculature, excluding singleton cells outside the body of the bud. Triplicate measurements were taken and averaged by an analyst blind to genotype. Tail myomeres were counted in ImageJ using the multipoint tool, starting at the somite directly dorsal to the vent and continuing through to the last distinct segment. To quantify *pah* and *nkx3-1* HCR staining, confocal z-stacks were imported into Imaris (Bitplane). Surfaces were created to define regions of interest (ROI) using absolute thresholding with a Surfaces Detail level of 2.4 (*pah*) or 2.41 (*nkx3-1*) and threshold values of 150 (*pah*) and 1100 (*nkx3-1*). The Volume Sum parameter was extracted from the final Surface for each sample and use for analysis in Prism 11 (GraphPad). For the *nkx3-1* ROI, all signal posterior to the fifth segment anterior to the end of the yolk extension was first captured. A second ROI was calculated if there was any signal below the body proper, then the volume of the second ROI was subtracted from the first to yield sclerotome volume. Datasets were tested for normality then analyzed with Student t-tests or non-parametric Mann-Whitney tests (2 samples) or Kruskal-Wallis tests with Dunn’s multiple comparison tests (3 samples) as indicated. Differences between genotypes were considered significant at *p*<0.05.

### scRNAseq protocol and analysis

*pax9^el622^* wild-type and mutant siblings were generated through an in-cross of heterozygotes, then subsequently in-crossed to produce clutches consisting entirely of wild-type or mutant embryos. At 3 dpf, healthy embryos were anesthetized, and the tail was removed with a micro-knife, cutting immediately anterior to the characteristic melanocyte gap. Seventy-five tails were pooled for the *pax9* mutant and wild-type samples, respectively, then dissociated into a single-cell suspension largely as described (Bresciani et al., 2018), with Liberase substituting for collagenase. Tail biopsies were washed in PBS, then incubated in a sterile dish with a pre-warmed dissociation mixture of Liberase and Trypsin-EDTA (40 μl/8% 25x Liberase: 460 μl Trypsin-EDTA) in a standard incubator set at 30°C with the plate placed directly on a heat block. Dissociation progress was monitored over 15-20 minutes, and gentle trituration was performed every 3 minutes with a P200 pipette tip against the bottom of the dish, being careful to avoid bubbles and aggressive pipetting. Once a single-cell suspension was achieved, a stop solution of Leibovitz’s media containing 10% FBS was added to halt dissociation. Pellets were washed in PBS containing 1% fetal bovine serum, filtered, and resuspended in 50 µl of 1% FBS/PBS. Samples were kept on ice and promptly submitted to the CCHMC Integrated Genomics and Microbiome Sequencing facility, where cell viability was confirmed at over 79%. The scRNA-seq assay was performed on a 10X Genomics Chromium X instrument, and libraries were prepared using the Chromium GEM-X 3’ Kit v4. Sequencing was performed on a NovaSeq X Plus Sequencer.

Raw sequencing data were processed into feature-barcode matrices using CellRanger v.9.0.1 in the 10x Genomics Cloud Analysis platform. The *Danio rerio* genome annotation release 106 was used for alignment (GRCz11). For the wild-type sample, 15,368 cells were captured, with 79% of reads mapping confidently to the genome (96.2% mapped overall) and an average of 13,231 reads/cell (mean) and 1301 genes per cell (median). For the mutant sample, 15,616 cells were captured, with 77.2% of reads mapping confidently to the genome (96.1% mapped overall) and 12,950 reads per cell and 1373 genes per cell. The data were then processed with single cell data analysis software Seurat (v.5.1.0). Seurat objects were created for each dataset in RStudio. Multiplets were detected using scDblFinder, run on each sample independently after a mild quality filter (nFeature_RNA>200 and percent.mt<6); the data were then filtered to save only singlets. After further high-stringency filtering (nFeature_RNA<4,500), 12,519 wild-type and 13,772 mutant cells were left. The wild-type and mutant Seurat objects were then merged for downstream analysis. No batch correction was applied because the samples were processed and sequenced in parallel, and we prioritized preservation of biologically meaningful differences between genotypes. Normalization was performed with the LogNormalize() method and a scale factor of 10,000. Informative genes were identified with FindVariableFeatures() using nfeatures=2,000. Following PCA, K-nearest neighbor graphs were constructed using FindNeighbors() with k=20 and dims=1:20, and clusters were determined with FindClusters() at a resolution of 0.25. Significantly enriched marker genes for each cluster were called with FindAllMarkers(), and cluster identity was determined by cross-referencing gene lists with the Daniocell scRNAseq database (Sur et al., 2023) and published in situs (www.zfin.org). Cell cycle phase coding was performed as a descriptive measure with the CellCycleScoring() function. Differential expression between mutant and wild-type cells was calculated with Wilcoxon rank tests or linear regression, as indicated in the text, and min.pct=0.1. To control for differences in detection rate between mutant and wild-type mesenchymal cells, nFeature_RNA and nCount_RNA were included as latent variables in the linear regression model. Volcano plots were generated with ggplot2. Those genes with a *p*-adj <0.05 and log2 fold-change threshold greater than 0.58 or less than -0.58 are shown as purple or blue colored dots in Figs. S4, S7, and S9, respectively. Gene expression across the entire dataset or the mesenchyme subcluster was visualized using the FeaturePlot(), DotPlot(), or VlnPlot() function. Microsoft Copilot was used to troubleshoot the R script.

## DATA AVAILABILITY

scRNAseq data are available at NCBI Gene Expression Omnibus under accession GSE328963. The R script is available from the corresponding author upon request.

## ACKNOWLEDGMENTS

We thank the fish facility staff at CCHMC, CU-AMC, Purdue University, and the University of Calgary for expert fish care; Kelly Rangel and the CCHMC Integrated Genomics and Microbiome Sequencing Facility for performing the scRNAseq assay; Krishna Roskin and the CCHMC IS4R group for bioinformatics assistance; and the CCHMC Bio-Imaging and Analysis Facility for microscope access and support. We thank Chunyue Yin for sharing the *hand2:EGFP* and *hand2^Cre^* lines. We are especially indebted to Ferenc Mueller, Matthew Harris, and Justin Cotney for insightful scientific discussions. This project was supported by the Cincinnati Children’s Research Foundation (lab start-up funds and a Research Innovation Pilot project to L.B.), the National Institutes of Health (2R35GM124913 to G.Z. (*pax9^pu115^* and *pax9^pu116^* mutant-related) and 3T32GM141742-03S1 to M.K.), the National Science Foundation (Graduate Research Fellowships Program 201569 to R.B.-Z.), and the Canadian Institutes of Health Research (project grants MOP-136926 and PJT-169113 to P.H., which covered all work performed at the University of Calgary (*pax9^ca122^*and *pax9^ca123^* mutant-related)).

## AUTHOR CONTRIBUTIONS

*Conceptualization* – L.B. & J.T.N.; *Investigation* – S.M., M.K., L.B., Z.D., R.B.-Z., K.M.K., S.P., A.M.-M., M.S.-P., C.A.H., & R.B.; *Formal analysis* – S.M., M.K., L.B., & K.M.K.; *Supervision*: L.B., J.T.N., G.Z., & P.H.; *Funding acquisition* – L.B., G.Z., P.H., J.T.N., M.K., & R.B.-Z.; *Writing (original draft)* – S.M. & L.B.; *Writing (review & editing)* – L.B., M.K., J.T.N., G.Z. & P.H.

