## Supplementary Figures S1-S9 for "Pax9 governs anterior identity and deployment of sclerotome to the median fins"

**Table S1. Enriched genes by cluster for full tail dataset**

**Table S2. Differential expression analysis of full tail dataset**

**Table S3. Enriched genes by cluster for mesenchyme subset**

**Table S4. Differential expression analysis of mesenchyme subset**

**Table S5. Differential expression analysis of *pax9*-expressing mesenchyme**

**Movie S1. The ectopic fin does not impede swimming in *pax9* mutant zebrafish**

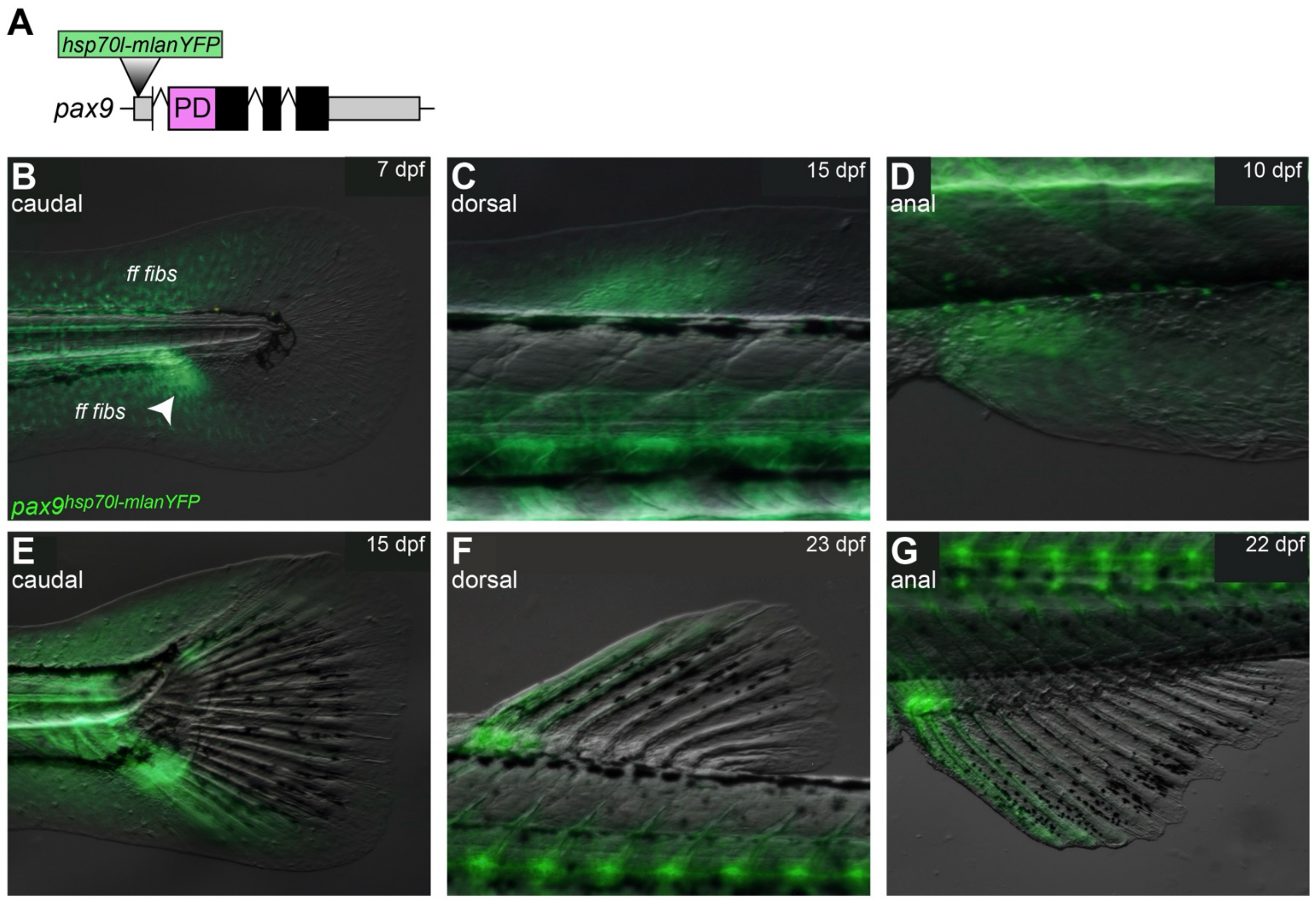


**Fig. S1. Early *pax9* knock-in reporter activity in median fins. A**, Location of targeted insertion in the *pax9* 5'UTR. **B-G**, Reporter activity in early buds and mid-development of the caudal (B, E), dorsal (C,F) and anal (D,G) fins. Arrowhead in B points to enriched activity in the anterior caudal bud. ff fibs, fin fold fibroblasts.

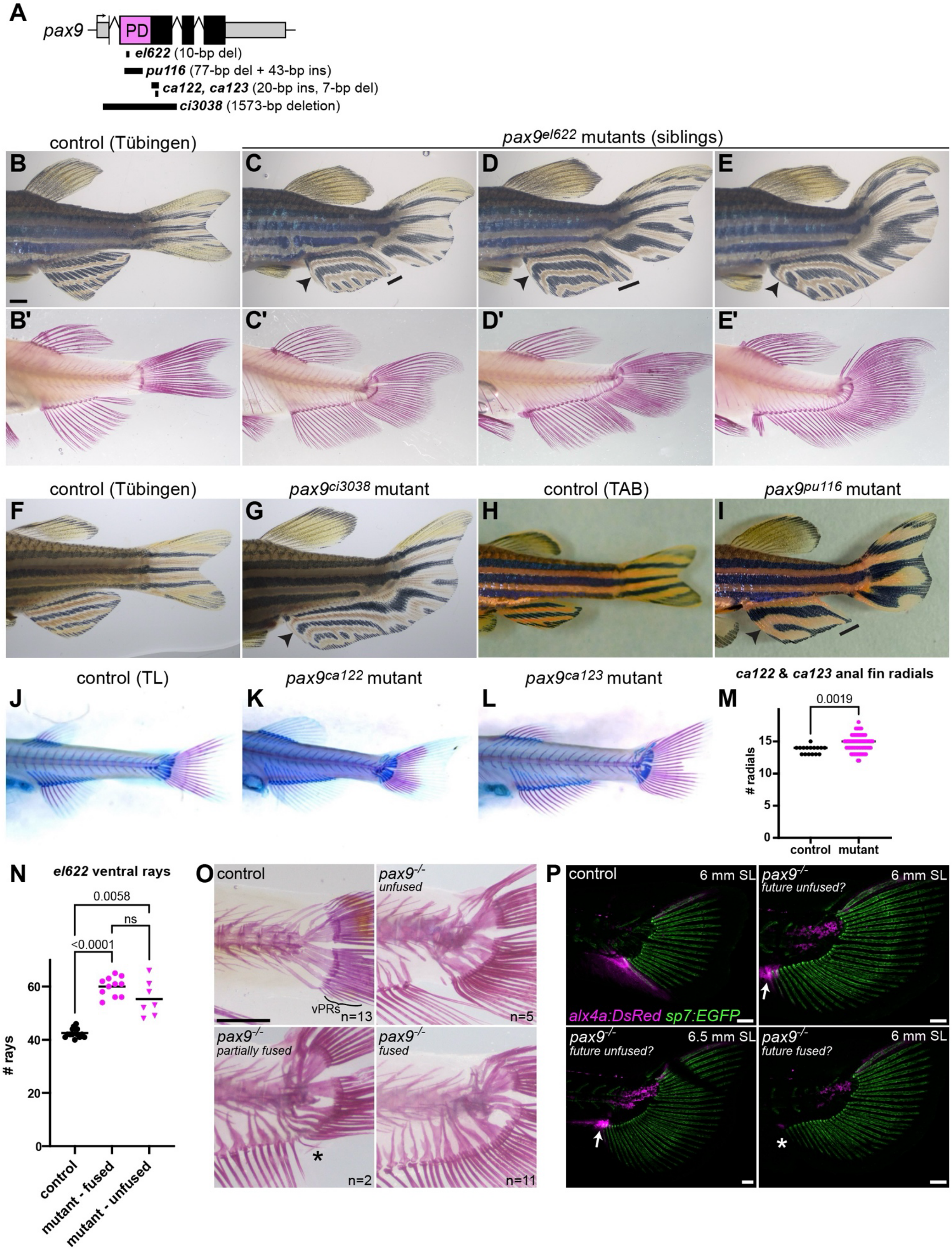

A
pax9
PD
el622 (10-bp del)
pu116 (77-bp del + 43-bp ins)
ca122, ca123 (20-bp ins, 7-bp del)
ci3038 (1573-bp deletion)
control (Tübingen)
pax9el622 mutants (siblings)
B
C
D
E
B'
C'
D'
E'
control (Tübingen)
pax9ci3038 mutant
control (TAB)
pax9pu116 mutant
F
G
H
I
control (TL)
pax9ca122 mutant
pax9ca123 mutant
ca122 & ca123 anal fin radials
J
K
L
M
0.0019
### radials
control
mutant
N
el622 ventral rays
0.0058
ns
<0.0001
### rays
control
mutant - fused
mutant - unfused
O
control
vPRs
n=13
pax9-/-
unfused
n=5
pax9-/-
partially fused
n=2
pax9-/-
fused
n=11
P
control
6 mm SL
alx4a:DsRed sp7:EGFP
pax9-/-
future unfused?
6 mm SL
pax9-/-
future unfused?
6.5 mm SL
pax9-/-
future fused?
6 mm SL

**Fig. S2. Variation in median fin expressivity across *pax9* mutant lines. A**, Approximate locations of the five *pax9* mutant alleles. PD, paired domain. **B-E**, Brightfield images of paired unprocessed and Alizarin red-stained *pax9*$^{el622}$ siblings, showing the phenotypic spectrum for this allele. The anal and caudal fins are fully fused in the specimen shown in E, partially fused in D, and unfused/discontinuous in C. Note that the anal fin is posteriorly elongated (black bars) and the caudal fin is asymmetric and anteriorly expanded in all. Arrowheads mark anomalies in anterior anal fin pigmentation. **F-L**, Four other independently generated *pax9* knockout alleles, *pax9*$^{ci3038}$ (F-G), *pax9*$^{pu116}$ (H-I), *pax9*$^{ca122}$, and *pax9*$^{ca123}$ (J-L), show similarly robust (*ci3038*) or milder (*pu116*, *ca122*, *ca123*) median fin phenotypes compared with *pax9*$^{el622}$. In the weaker lines, the caudal fin remains asymmetric, and the anal fin is still elongated posteriorly (black bar in I) and aberrantly pigmented (arrowheads). Anal fin proximal radials for *ca122* and *ca123* quantified in M; significance determined by a Mann-Whitney test. **N**, *pax9*$^{el622}$ mutants with fully fused fins tend to have more rays than unfused/partially fused mutants but the difference was not significant (Kruskal-Wallis test with Dunn's multiple comparisons). **O**, Higher magnification images of the variable *pax9*$^{el622}$ phenotypes observed in the fusion region. vPRs, ventral procurrent rays. Asterisk marks the lone short procurrent ray in a partially fused mutant. **P**, A small patch of anterior caudal *alx4a:*DsRed signal is present in some juvenile *pax9*$^{el622}$ mutants (white arrows). This may correlate with later formation of a partially fused or discontinuous fin, versus complete absence of expression (bottom right, asterisk) presaging a fully fused fin. Images are maximum intensity projections. Scale bars in B-E, O = 1 mm, P = 100 µm.

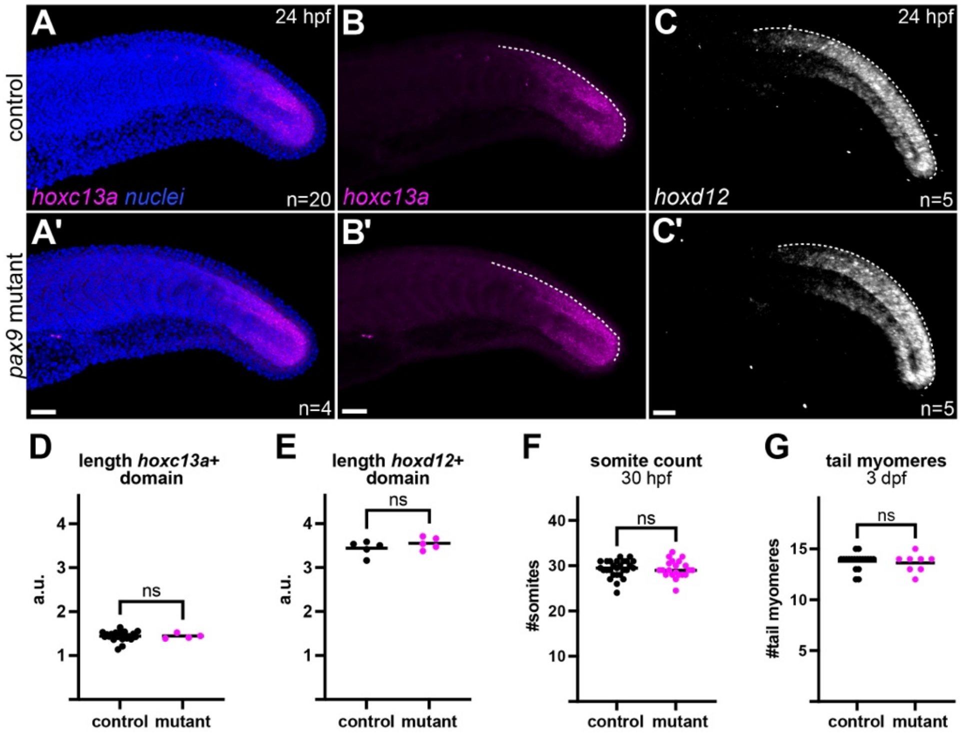


**Fig. S3. Loss of *pax9* does not impact global axial patterning. A-C**, HCR for *hoxc13a* (A-B) and *hoxd12* in 24 hpf control and *pax9* mutant embryos. The dotted line marks the dorsal margin of the hox expression domain. Nuclei are shown for context in A-A'. Scale bars = 50 µm. **D-E**, Length of the dorsal margin of the *hoxc13a*+ (D) and *hoxd12*+ (E) domains was similar across genotypes ($p$=0.7941, $p$=0.4206, respectively). **F**, Somite numbers, as counted by eye under transmitted illumination at 30 hpf, were unchanged between genotypes ($p$=0.6134). **G**, Tail myomere numbers in phalloidin-stained 3 dpf larvae were similar across genotypes ($p$=0.6068). Images are maximum intensity projections. Mann Whitney tests used for all comparisons ($p$>0.05).

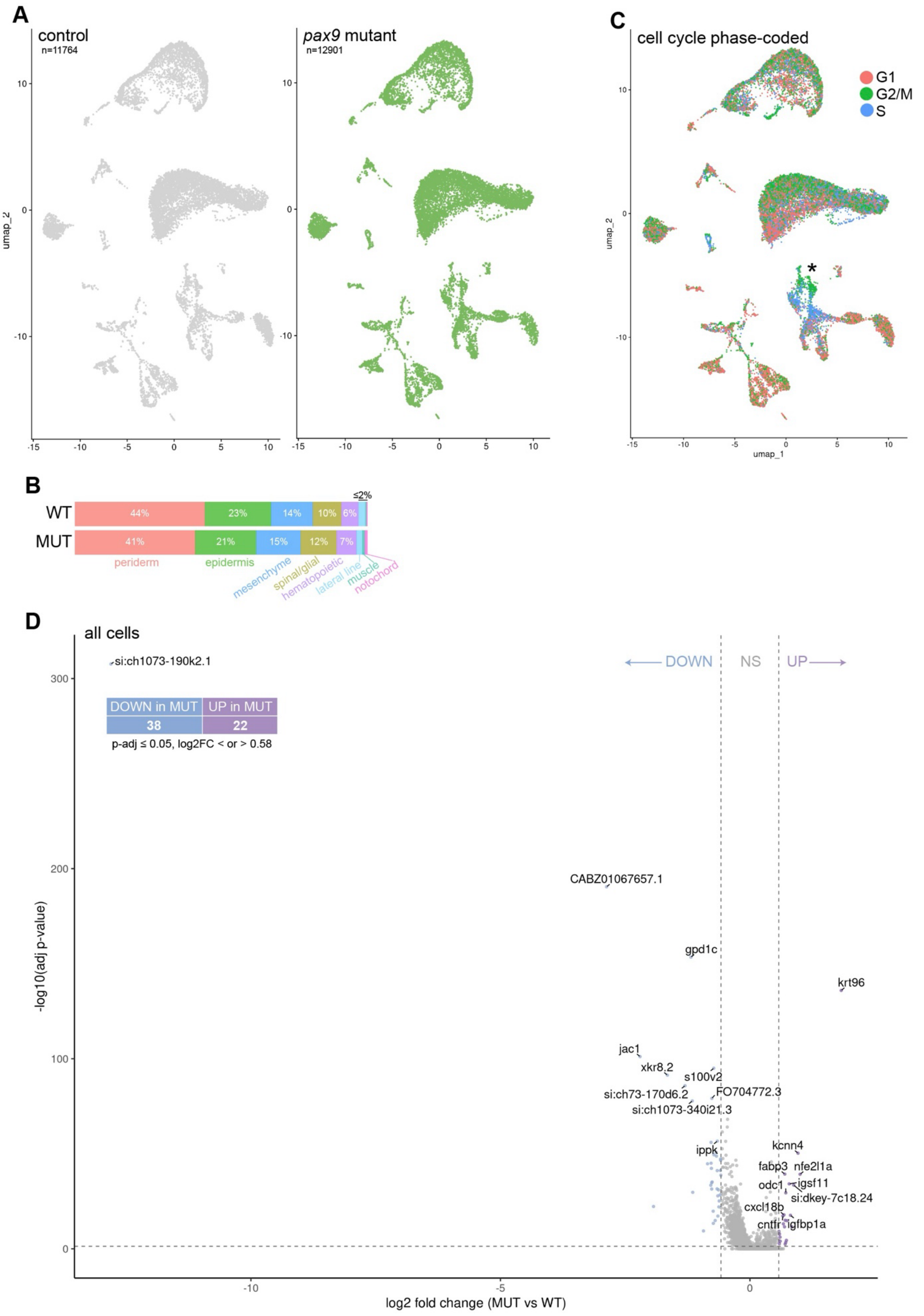

A
control
n=11764
pax9 mutant
n=12901
C
cell cycle phase-coded
G1
G2/M
S
umap_2
umap_1
B
WT
MUT
44%
23%
14%
10%
6%
≤2%
41%
21%
15%
12%
7%
periderm
epidermis
mesenchyme
spinal/glial
hematopoietic
lateral line
muscle
notochord
D
all cells
DOWN
NS
UP
DOWN in MUT
UP in MUT
38
22
p-adj ≤ 0.05, log2FC < or > 0.58
si:ch1073-190k2.1
CABZ01067657.1
gpd1c
krt96
jac1
xkr8.2
s100v2
si:ch73-170d6.2
FO704772.3
si:ch1073-340i21.3
ippk
kcnn4
fabp3
nfe2l1a
odc1
igsf11
si:dkey-7c18.24
cxcl18b
cntfr
igfbp1a
-log10(adj p-value)
log2 fold change (MUT vs WT)

**Fig. S4. Similarity of caudal fin scRNAseq samples from wild-type and mutant fish. A**, UMAPs for full dataset segregated by genotype. **B**, Proportions of each sample that fell into each major cell lineage. Periderm, epidermis, lateral line, hematopoietic, and spinal/glial subclusters were pooled for simplicity. **C**, UMAP of tail cells coded by cell cycle phase. Note most populations are not primarily defined by cell cycle, with the exception of the proliferating mesenchyme (asterisk). **D**, Volcano plot showing minimal differential expressed genes across the full dataset. The top ten most significantly up or downregulated genes are labeled.

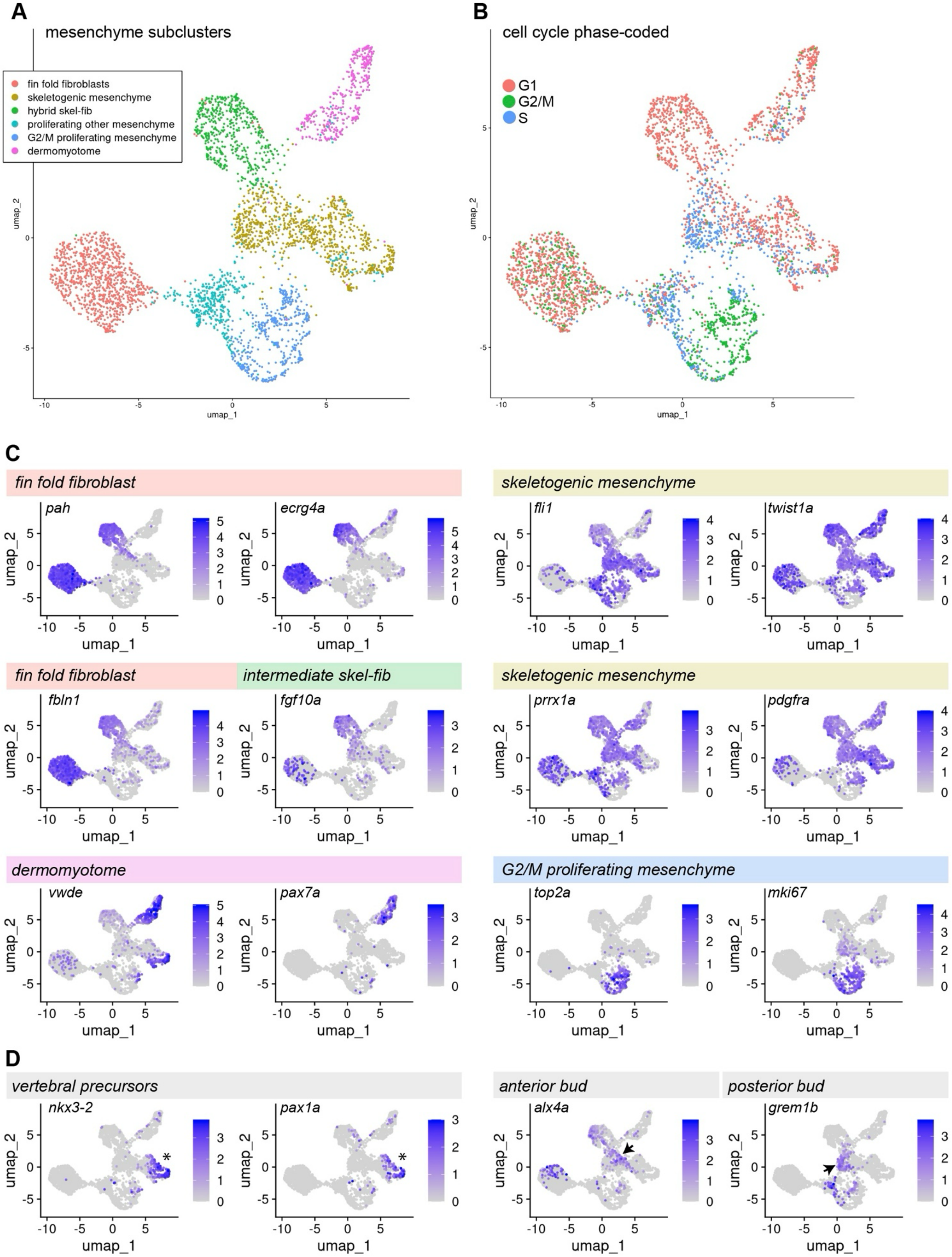

A
mesenchyme subclusters
fin fold fibroblasts
skeletogenic mesenchyme
hybrid skel-fib
proliferating other mesenchyme
G2/M proliferating mesenchyme
dermomyotome
umap_1
umap_2
B
cell cycle phase-coded
G1
G2/M
S
C
fin fold fibroblast
pah
ecrg4a
skeletogenic mesenchyme
fli1
twist1a
fin fold fibroblast
intermediate skel-fib
fbln1
fgf10a
skeletogenic mesenchyme
prrx1a
pdgfra
dermomyotome
vwde
pax7a
G2/M proliferating mesenchyme
top2a
mki67
D
vertebral precursors
nkx3-2
pax1a
anterior bud
alx4a
posterior bud
grem1b

**Fig. S5. Properties of mesenchymal subclusters. A**, UMAP of the mesenchyme/fibroblast clusters with both genotypes pooled (compare with Fig. 6C). **B**, UMAP of mesenchymal cells coded by cell cycle phase. **C**, FeaturePlots for marker genes used to determine cluster identity; cells from both genotypes are pooled. **D**, FeaturePlots for genes enriched in subregions of the skeletogenic mesenchyme cluster, predicted to represent vertebral precursors (asterisks) vs. anterior and posterior caudal fin bud cells (arrows).

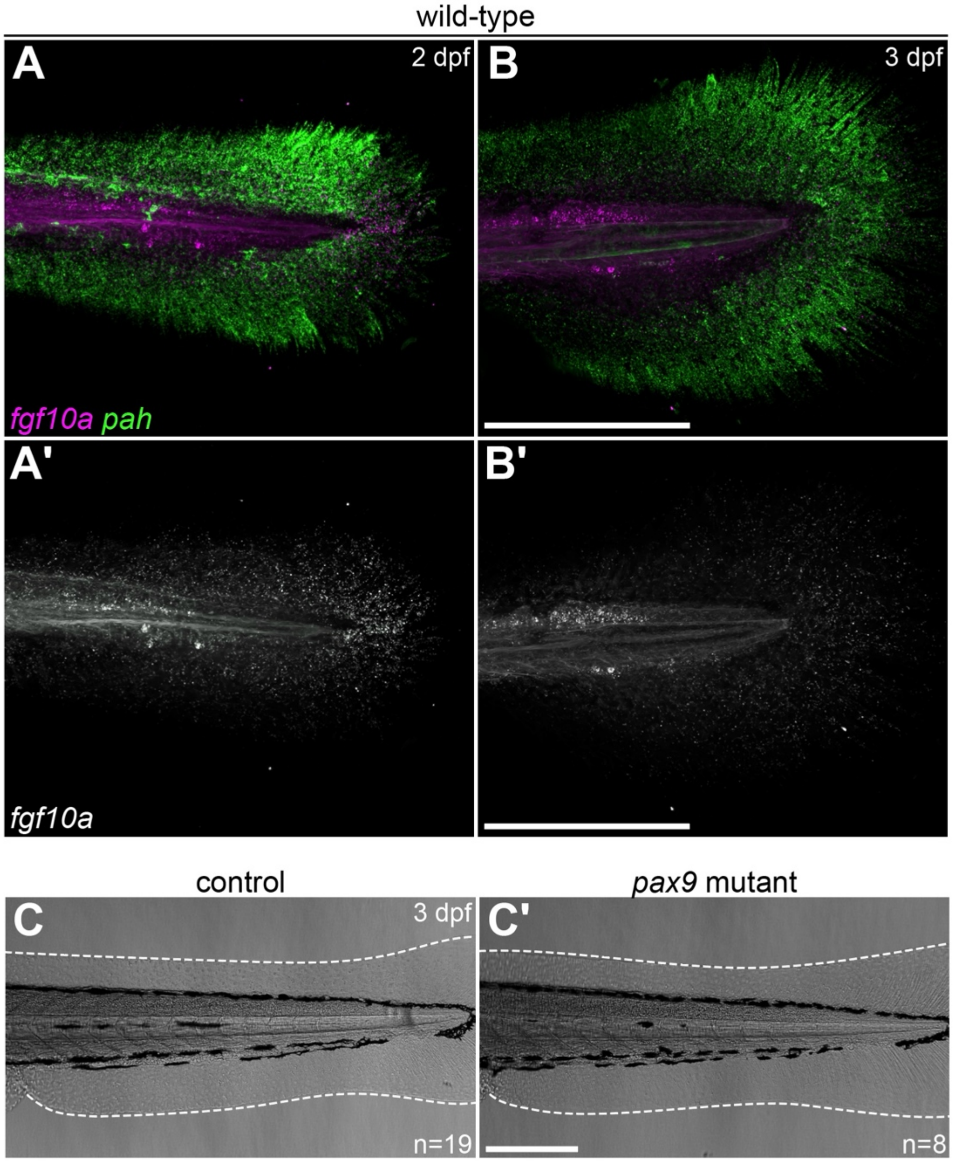


**Fig. S6. Fin fold features. A-B**, Compared with the broadly expressed fin fold fibroblast marker *pah*, *fgf10a* is restricted to the caudal tip of the fin fold of wild-type fish at 2 (A) and 3 dpf (B). These cells may be the final remnants of the *fgf10a*+ tail bud. Maximum intensity projections of denoised z-stacks, HCR. **C**, Transmitted light images of the tail region (vent to tip of notochord) in fixed 3 dpf larvae. The fin fold is not morphologically abnormal in mutants. Dashed line traces the edge of the fin fold. Scale bars in A-C = 250 μm.

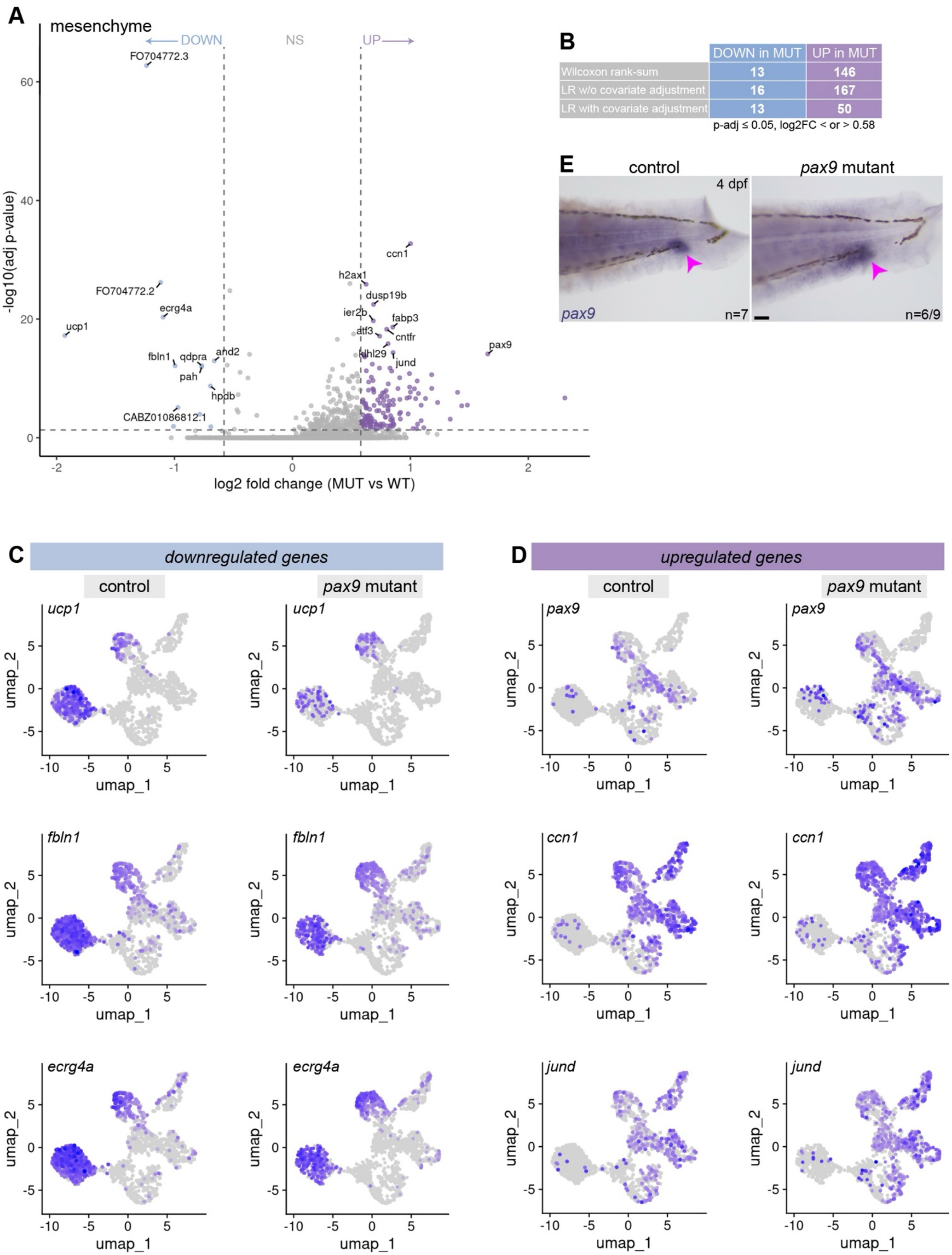


| | DOWN in MUT | UP in MUT |
|---|---|---|
| Wilcoxon rank-sum | 13 | 146 |
| LR w/o covariate adjustment | 16 | 167 |
| LR with covariate adjustment | 13 | 50 |

p-adj ≤ 0.05, log2FC < or > 0.58

**Fig. S7. Differential gene expression in the mesenchyme cluster supports reduction of fin fold fibroblasts. A**, Volcano plot for differentially expressed genes in the mesenchyme cluster (Wilcoxon analysis). **B**, The elevated number of upregulated genes is in part a technical artifact of higher detection rates in mutant cells, supported by a complementary linear regression analysis that used nFeatures_RNA and nCounts_RNA as covariates (B). **C**, Downregulated genes are enriched in fin fold fibroblasts and the hybrid population (see Fig. S5A for reference UMAP). **D**, Upregulated genes, including *pax9*, are more broadly expressed throughout the rest of the mesenchyme. **E**, Validation of upregulated *pax9* expression in the mutant tail (pink arrowhead). Scale bar = 50 μm.

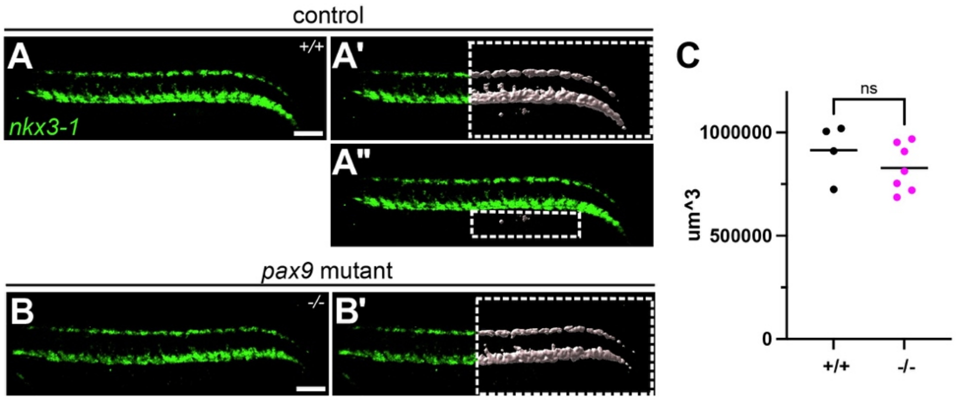


**Fig. S8. No overt changes in sclerotome volume in *pax9* mutants. A-C,** Sclerotome volume was measured in Imaris on samples processed for *nkx3-1* HCR (A). All signal posterior to the fifth segment anterior to the end of the yolk extension was first captured (A', B'), then a second ROI was calculated if there was any signal below the body proper (A''), then the volume of the second ROI was subtracted from the first to yield sclerotome volume. Volumes are graphed in C, with significance determined by a Mann-Whitney test. Images are maximum intensity projections.

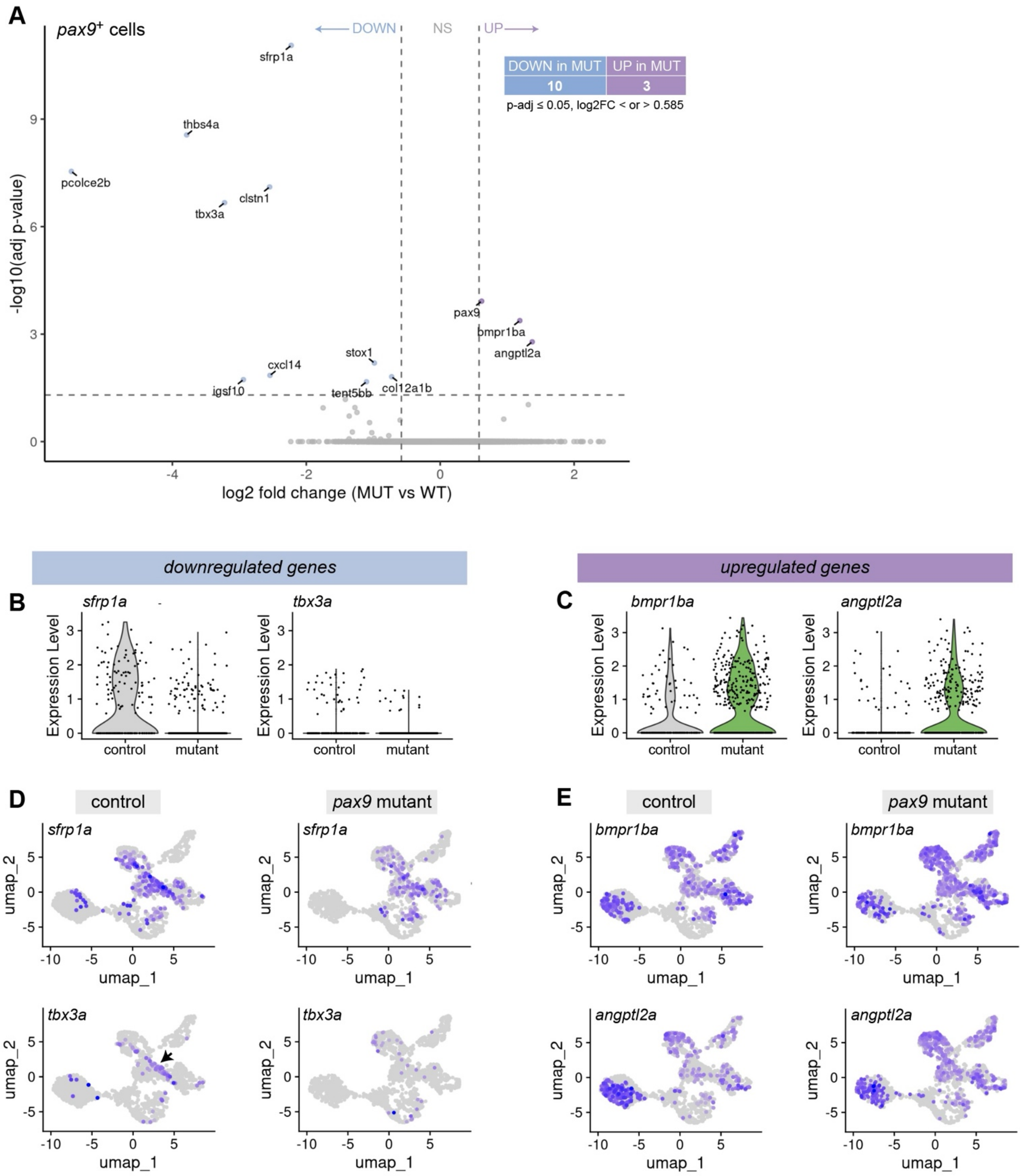


**Fig. S9. Differential gene expression analysis within *pax9*+ cells identifies candidate targets. A**, Volcano plot of differentially expressed genes in *pax9*-expressing cells. **B-C**, Violin plots for notable down- (B) and upregulated (C) genes within the *pax9*+ subset. **D-E**, Feature plots for the same dysregulated genes within the full mesenchyme dataset, split out by genotype. *tbx3a* in particular is enriched in the putative anterior bud domain in controls (arrow, compare with *alx4a* in Fig. S5D) but absent in mutants.
